# Novel mechanisms for phosphate acquisition in abundant rhizosphere-dwelling *Bacteroidetes*

**DOI:** 10.1101/719427

**Authors:** Ian D.E.A. Lidbury, David J. Scanlan, Andrew R. J. Murphy, Andrew Bottrill, Alex Jones, Mark Tibbett, Gary Bending, John P. Hammond, Elizabeth M. H. Wellington

## Abstract

Global food production is reliant on the application of finite phosphorus (P) fertilisers. Numerous negative consequences associated with intensive P fertilisation have resulted in a high demand to find alternative sustainable methods that will enhance crop P uptake. *Bacteroidetes*, primarily from the genus *Flavobacterium*, have recently been shown to be abundant members of the plant microbiome, but their general ecological role and potential to mobilise P in the rhizosphere remains very poorly characterised. Here, we sought to determine the P mobilisation potential of *Flavobacterium* strains isolated from the rhizosphere of oilseed rape (*Brassica napus* L.). We show that these *Flavobacterium* strains possess novel mechanisms for P mobilisation and subsequent acquisition. These include the constitutive and inducible expression of completely novel and phylogenetically distinct phosphatases, the phosphate starvation inducible expression of uncharacterised and hypothetical genes and gene clusters and, for the first time, the involvement of outer membrane SusCD transport complexes (usually associated with carbohydrate transport) in P acquisition. The genes encoding these unusual phosphate starvation inducible proteins were enriched in plant-associated *Flavobacterium* strains suggesting that this machinery represents niche-adaptive strategies for overcoming P scarcity in this genus. We propose that abundant rhizosphere-dwelling *Flavobacterium* spp. have evolved unique mechanisms for coping with Pi-stress which may provide novel solutions for future sustainable agricultural practices.

## Introduction

For all living organisms on earth, phosphorus (P) plays an integral role in maintaining normal cellular function due to its incorporation into several cellular components, such as nucleic acids, phospholipids and adenosine triphosphate (ATP). P is often a limiting nutrient for primary production in both terrestrial and marine biomes and therefore has a vital role in both terrestrial and marine carbon cycles, as well as global food security ^1–3^. Over the past 60 years agricultural systems have seen a tremendous rise in the use of phosphate (Pi) fertilisers derived from rock phosphate, with current estimates showing we apply over 20 million tons year^−1^ to sustain adequate yields ^1^. Rock phosphate is a finite commodity and there are major concerns regarding the long-term availability of this valuable resource in global food production ^1,4^. Furthermore, on average only 20% of the phosphate applied is consumed by humans and the majority of excess P is either retained by soils or dissipates into neighbouring ecosystems with detrimental effects on biodiversity^5^. In addition, the availability and mobilisation of P has also been linked to the productivity of tropical forests and their ability to sequester atmospheric CO_2_ (Goll 2012; Yang et al., 2016). Therefore, a greater understanding of the soil microbial P cycle will have a number of important consequences for climate change models and global food security. Organic P (P_o_) often accounts for the majority of total P in soils and must be solubilised by soil microorganisms before becoming available for uptake by plants^4,6^. Despite this, research and practices tend to focus on increasing the availability of Pi and often neglect the P_o_ cycle ^7^.

Bacteria have evolved numerous strategies to overcome environmental Pi scarcity which are almost exclusively regulated by a two-component regulatory system, usually referred to as PhoBR (PhoB, DNA binding response regulator; PhoR, transmembrane histidine kinase) ^8^. Several P scavenging strategies are common to almost all bacteria studied to date including: (1) the production of extracellular enzymes to hydrolyse Pi from various organic complexes (2) the release of organic acids through the catabolism of sugars (3) a reduction in the quota of phospholipids constituting their plasma membranes (4) up-regulation of transporters for acquiring simple organic P forms, e.g. phosphonates or glycerol-3-phopshate (5) up-regulation of the high affinity phosphate transport system, commonly known as PstSABC ^8–13^. Several studies have shown that Bacteria predominantly secrete non-specific alkaline phosphatases (APase, three distinct forms known as PhoA, PhoD and PhoX) whilst a few cultured strains secrete acid phosphatases (AcPases, class I, II and III) in response to Pi limitation ^13–17^. The original paradigm was that all Bacteria possess one of these enzymes, but recent evidence from genomic and metagenomic datasets suggest that only half possess these P mobilising enzymes ^18^. In a similar manner it is generally accepted that all Bacteria capable of responding to Pi limitation possess at least one form of PstSABC and that this interacts with PhoBR to help regulate the Pi stress response^8^.

*Flavobacteriaceae* and *Sphingobacteraceae* belong to the phylum *Bacteroidetes* and are dominant members of plant, soil and ocean microbiomes^19–25^. In ocean microbiomes, they are crucial members of the community associated with the degradation of algal-derived polysaccharides and are thus key regulators of the global ocean carbon cycle ^23,26^. Terrestrial *Flavobacteriaceae* have also been associated with the breakdown of plant-derived glycans due to the high number of hydrolytic enzymes encoded in their genomes ^27,28^. Another molecular signature associated with polysaccharide degradation is the presence of a two-component outer membrane transport system, termed SusCD (archetypal form, **<u>S</u>**tarch **<u>u</u>**tilisation **<u>s</u>**ystem). SusC is a transmembrane TonB-dependent transporter (TBDT) and SusD is the ligand-binding lipoprotein cap^29^. SusCD complexes are usually constituents of specialised operons, termed polysaccharide utilisation loci (PUL). The specificity of the SusCD-complexes allows *Bacteroidetes* to grow on a variety of polysaccharides ^30,31^ and these transporters are used for predicting the *in situ* degradation of plant or algal-derived polysaccharides ^32^.

In the rhizosphere, *Flavobacteriaceae* typically account for 5-65% of the microbial community associated with various agriculturally important crops ^20,21,24,33–35^. Indeed, *Flavobacteriaceae* abundance in the rhizosphere is usually significantly higher than that of the surrounding bulk soil, suggesting that the plant microbiome provides a distinct niche for *Flavobacteriaceae* to occupy^20,24,33^. *Flavobacteriaceae* have been shown to play a role in defence against plant pathogens and can provide beneficial effects on plant growth, including P solubilisation, the production of plant growth hormones and antimicrobial production ^36–41^. In response to the realisation that *Flavobacteriaceae* are abundant in plant microbiomes, successful efforts have been made to develop suitable media for their cultivation ^42^. However, unlike *Pseudomonas*, *Bacillus* and *Streptomyces*, *Flavobacteriaceae* have not been extensively screened for their phosphate mobilisation capabilities. This is despite the fact that 25% of the isolates obtained from the Barley rhizosphere were related to *Flavobacteriaceae* (compared to less than 5% for *Pseudomonas*) and these contributed almost 50% of the potential microbial P-turnover ^33^.

In this study, we utilised a newly developed medium^42^ to isolate novel strains from the rhizosphere of the economically important crop *Brassica napus* L. (oilseed rape, OSR), grown under field conditions. The aim of the study was to identify the genes and enzymes expressed under Pi depletion in laboratory conditions to determine the phosphate-stress response (PHO) regulon of this understudied family. This would enable the potential identification of novel enzymes with biotechnological capability as well as identify biomarkers for future metaproteomic studies *in situ*. Our results revealed that *Flavobacteriaceae* exhibit unusual phosphatase activity, express a novel suite of Pi-responsive enzymes and also use novel mechanisms for Pi transport.

## Results

### Phenotypic and genomic characterisation of *Flavobacterium* OSR isolates

Initial screening for APase activity in *Flavobacterium* strains revealed that *F. johnsoniae* DSM2064, *Flavobacterium* sp. F52 (F52) and *Flavobacterium* sp. L0A5 (LOA5) all exhibited unusual constitutive APase activity during growth under both Pi replete and Pi deplete conditions (Figure 1A). To determine whether this was a more general trait of rhizosphere-associated *Bacteroidetes*, numerous strains were isolated from the rhizosphere of field-grown *B. napus*, using a semi-selective medium targeting *Bacteroidetes*^42^ and screened for their phosphatase activity under both Pi-replete and Pi deplete growth conditions. The majority of strains isolated were related to *Flavobacterium*. However, we also isolated *Chryseobacterium* and *Sphingobacterium*, as well as strains not related to *Bacteroidetes,* non-*Bacteroidetes* (NB), predominantly related to *Xanthomonas* and *Pseudomonas* (Figure S1). When grown on glucose, all strains related to *Bacteroidetes* exhibited constitutive APase activity under Pi-replete conditions, whilst only one out of the 14 NB rhizosphere isolates exhibited constitutive APase activity (Figure 1B). The majority of the *Bacteroidetes* strains failed to grow on succinate as a sole carbon source, but those that did also exhibited constitutive APase activity (Figure 1B). In contrast, no NB isolates exhibited APase activity when grown under Pi-replete conditions with succinate as a sole carbon source. All *Bacteroidetes* and NB isolates that grew exhibited APase activity when grown under Pi-deplete conditions. Importantly, these *Flavobacterium* strains showed superior APase activity during Pi-deplete growth on glucose (which simulates lower pH conditions more akin to the rhizosphere) compared to *Pseudomonas* rhizosphere strains^13^ whose activity almost disappeared compared to growth on succinate (Figure 1).

**Figure 1.**
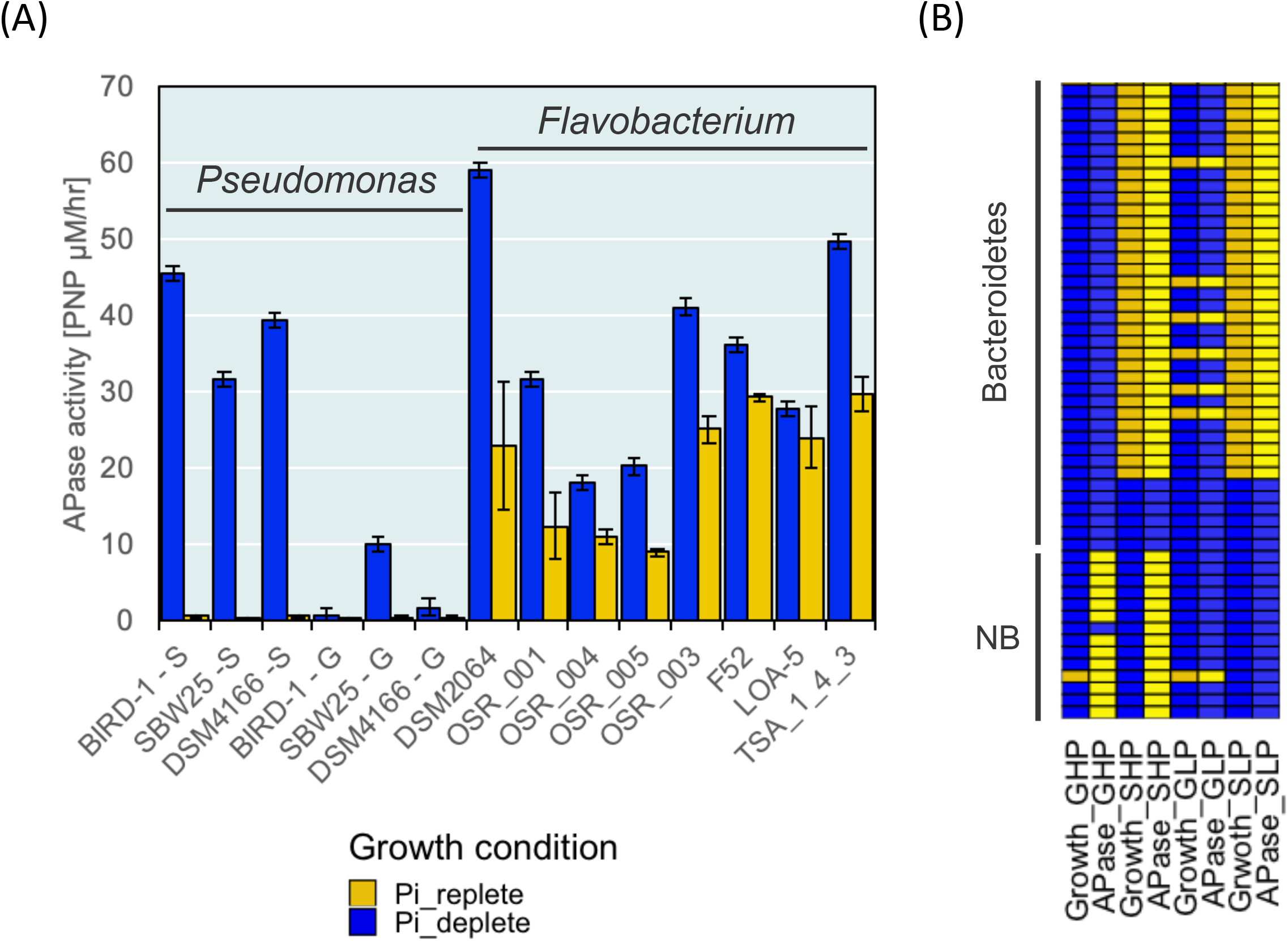
**(A).** Phosphatase activity of selected *Flavobacterium* strains including the oilseed rape (OSR) isolates. *Pseudomonas putida* BIRD-1, *Pseudomonas fluorescens* SBW25 and *Pseudomonas stutzeri* DSM4166 are also included as soil bacteria typically showing inducible phosphatase activity. Abbreviations: S, succinate-grown; G, glucose-grown. Note that *Flavobacterium* does not grow on succinate as a sole carbon source. Phosphatase activities were calculated in triplicate and error bars denote standard deviation. (**B).** Phosphatase activity was also recorded (n=3) for various *Bacteroidetes* and non-*Bacteroidetes* (NB) isolates grown on either glucose (20mM) or succinate (20mM) under Pi-replete or Pi-deplete conditions. Blue represents growth and positive activity and orange denotes lack of growth and yellow denotes lack of activity. Abbreviations: GHP, glucose high Pi; GLP, glucose low Pi; SHP, succinate high Pi; SLP, succinate low Pi.

Six isolates related to *Flavobacterium*, all exhibiting constitutive phosphatase activity (Figure 1A), were chosen for genome sequencing and subsequent exoproteomic analysis in addition to the other three *Flavobacterium* strains (*F. johnsoniae* DSM2064, F52 and LOA5) whose genomes were already sequenced. The newly isolated strains, referred to as OSR001/2/3/4/5 and TSA_1_4_3, captured the breadth of terrestrial *Flavobacterium* diversity ^27^ (Figure 2A). Whole genome alignment of the newly isolated *Flavobacterium* strains, as well as four genomes (deposited in IMG/JGI) from other root-associated *Flavobacterium* spp. and F52, possessed more streamlined genomes than *F. johnsoniae* DSM2064 (Figure 2B). TSA_1_4_3 had the largest genome (6.1 Mb) and OSR_004 had the smallest (4 Mb). Details of their general genomic characteristics are given in Table S1. Genome completeness was confirmed by screening for the presence of ∼80 core genes (Figure 6A). All *Flavobacterium* OSR isolates possessed homologs for the genes encoding the two-component master regulator PhoBR (Table S2). Unfortunately, after numerous attempts, we failed to generate a *phoBR* knock out mutant in *F. johnsoniae* DSM2064, suggesting that in this bacterium *phoBR* is essential for survival and precluding mutagenesis as a route to define the PHO regulon in this instance. Interestingly though, the *Flavobacterium* OSR isolates lacked numerous well-known genes associated with the PHO regulon. For example, only one newly isolated strain, OSR004 possessed genes (*pstSABC*) encoding the high affinity Pi transporter, PstSABC, though all nine strains possessed a gene predicted to encode the low-affinity Pi transporter PitA. All nine strains lacked PhoD, typically one of the most abundant APases found in soils and soil-dwelling bacteria^43^. *F. johnsoniae* DSM2064, F52, LOA-5 and four out of the six OSR isolates, including TSA_1_4_3, possessed one or two genes with homology to PhoA (pfam00245) though the identity of the translated amino acid sequences was low (~25% identity to *E. coli* PhoA, Figure S1). Furthermore, BLASTP (using either *P. putida* BIRD-1 or *Ruegeria pomeroyi* DSS-3 as queries) or Pfam searches only detected the presence of PhoX in TSA_1_4_3, the APase responsible for the vast majority of phosphatase activity in rhizosphere-dwelling *Pseudomonas* and marine *Roseobacter* strains^13,44,45^. In contrast, all strains possessed a homolog of the gene encoding the beta-propeller phytase within their genomes (Table S2).

**Figure 2.**
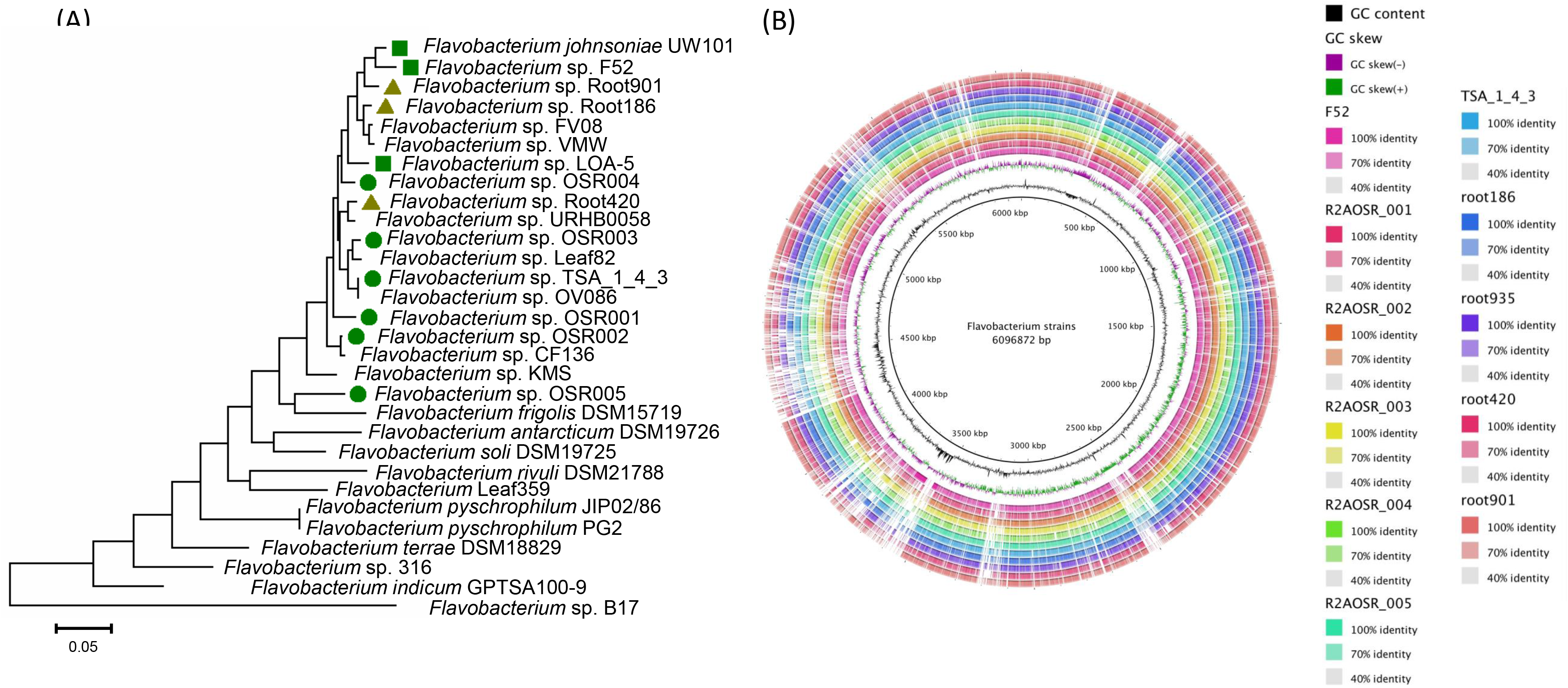
**(A)** Phylogenetic analysis of *Flavobacterium* isolates using a multi-locus approach with 10 single copy core genes (as previously described by Kolton et al. 2013). Evolutionary distances were inferred using the maximum-likelihood method. The *Brassica napus* isolates (green circles) spread the breadth of terrestrial *Flavobacterium* strains and captured more diversity than DSM2064 and Israeli strains tested (green squares) or the *Arabidopsis* strains (olive green triangles). (**B)** BRIG analysis of the oilseed rape (OSR) isolates, F52, and four root-associated *Flavobacterium* strains compared against *F. johnsoniae* DSM2064 revealed common patterns of genomic architecture among the strains.

### Proteomic response linked to Pi mobilisation

To determine the proteomic response of *Flavobacterium* strains to Pi-depletion, the nine *Flavobacterium* strains were subjected to growth under Pi replete or Pi deplete conditions. Unfortunately, for OSR002 we were unable to adequately extract exoproteins due to this culture’s inability to form a coherent cell pellet following centrifugation. Based on their exoproteomes, all eight strains elicited a clear proteomic response to Pi depletion (Figure 3, Table S3-S10). Interestingly, we identified numerous proteins that were either very distinct compared to previously characterised Pi acquisition proteins^8,13,14^, were completely novel proteins of unknown function, or were novel Pi-stress response proteins containing Pfam domains linked to broad protein families. The major findings from these exoproteomics data can be categorised into three major categories described below.

**Figure 3.**
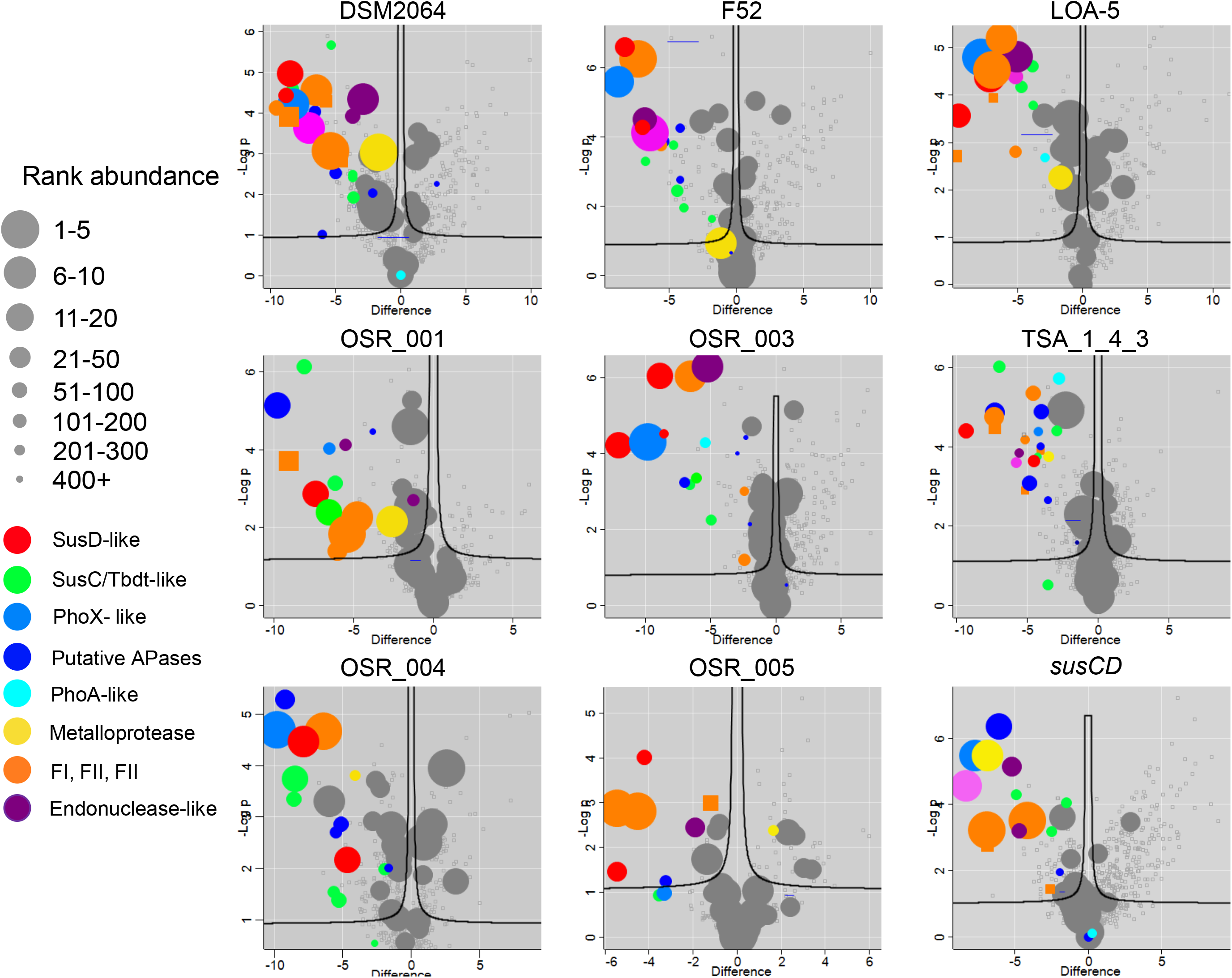
Exoproteomic profiles of *Flavobacterium* strains grown under Pi-replete or Pi-deplete conditions. Volcano plots illustrate both the difference in the log2 label free quantification (LFQ) intensity between treatments and the log probability of the observed difference. Negative difference values represent proteins enriched in Pi-deplete compared to Pi-replete and vice versa. The rank abundance is denoted by dot size. All of the top 30 proteins are highlighted. After this, only the rank abundance of proteins showing a significant difference or ones that may be linked to the PHO regulon are highlighted. All experiments were performed in triplicate and mean values have been plotted. Abbreviations: PusD, phosphate utilisation system-like outer membrane binding domain; PhoX, alkaline phosphatase PhoX-like; SusC/TonB-like, including TonB-dependent transporter-like proteins; APases, putative phosphatases; FI, FII, FIII, refer to the hypothetical exoproteins that share weak homology with pectate lyases/polygalacturonases.

### *Flavobacterium* strains express numerous uncharacterised proteins with potential APase activity

Under Pi depletion, all *Flavobacterium* strains expressed and secreted numerous uncharacterised proteins containing Pfam domains Pfam00149, Pfam04231, Pfam01676, Pfam01663, Pfam03372 associated with various phosphomonoesterases and phosphodiesterases (Figure 3, blue and maroon circles) that are distinct from the classical PhoX (Pfam05787), PhoD (Pfam09423/Pfam16655), UshA (Pfam02872) and GlpQ (Pfam03009) proteins typically associated with soil bacteria. Surprisingly, whilst some of the isolates (TSA_1_4_3, LOA-5, OSR001 and OSR003) secreted PhoA-like proteins (Pfam00245) during Pi-depletion, these predicted APases did not form a major component of the Pi-deplete *Flavobacterium* exoproteomes (Table S4-S10). In particular, in DSM2064, PhoA1 (Fjoh_3249) was not detected and only a few peptides belonging to PhoA2 were detected in two Pi-deplete replicate cultures. However, all *Flavobacterium* strains did secrete a predicted lipoprotein in response to Pi-depletion that was frequently among the top five most abundant and the most differentially expressed exoproteins (Figure 3, Tables S3—S10). Further bioinformatics analysis revealed this protein has a region with weak homology to DUF849 (Expected value ~−6^−11^), that is typically associated with PhoX. However, the *Flavobacterium* PhoX-like proteins are phylogenetically distinct compared to PhoX found in other taxa, such as *Proteobacteria*, *Actinobacteria* and other *Bacteroidetes* (Figure S3). In addition, all strains also up-regulated one or two lipoproteins (Fjoh_4721 and Fjoh_4723 in DSM2064) containing the Pfam04321 domain found in the characterised periplasmic endonuclease_1 (EndA) of *E. coli*, *Vibrio cholerae* and *Aeromonas hydrophila*^46^. The *Flavobacterium* EndA-like exoproteins are over twice the length of *Gammaproteobacteria*-related EndA homologs and are phylogenetically distinct, likely due to differences in their cellular localisation (Figure S4). In a few strains, some of these phosphodiesterases and phoshomonoesterases were co-localised in low Pi-responsive clusters, which also contained TBDTs, sugar P permeases, a novel P-related transporter (discussed in the next section), predicted intracellular and extracellular nucleases, oxidoreductases, metal-dependent hydrolases and a putative phosphonate hydrolase (PhnX) (Figure S5). Seven of the nine strains also expressed a lipoprotein containing a predicted endo/exonuclease/phosphatase domain (Pfam_03372). This exoprotein (Fjoh_0074 in DSM2064, Table S2-9) was expressed in both conditions and very abundant in some isolates (F52) and is therefore a candidate for the constitutive APase activity.

### *Flavobacterium* strains express uncharacterised outer membrane SusCD complexes in response to Pi depletion

Whilst the majority of expressed SusCD complexes, including previously characterised forms^47^, showed no change in expression in response to Pi concentration (Figure 4A), all *Flavobacterium* isolates induced the expression of certain SusCD-like complexes in response to Pi depletion (Figure 4A). Furthermore, all isolates, whose genomes lacked the high affinity Pi transporter PstSABC (7/8), abundantly expressed one or two particular forms of SusCD, hereafter termed **<u>P</u>**hosphate **<u>u</u>**tilisation **<u>s</u>**ystem (Pus)CD1 and PusCD2 (Figure 4A and B). The genes (*pusCD1* and *pusCD2*) encoding PusCD1 (Fjoh_4168 in DSM2064) and PusCD2 (Fjoh_3250) were co-localised with various phosphatase/esterase-like genes, including *phoA1* (Figure 4C). *pusCD1* and *pusCD2,* were not found in the genome of OSR_004, which did possess *pstSABC*. Searching 100 *Flavobacterium* genomes deposited in the Integrated Microbial Genomes database (Table S12) further confirmed that the presence of *pusCD1*, and to a lesser extent *pusCD2*, was coincident with a lack of *pstSABC* (Figure 4D). *pusD3* represented a distinct form of *pusD1* (Figure 4B), but had a similar genetic neighbourhood to that of *pusD2. pusD4*, *pusD5* and *pusD6*, were all co-localised with novel clusters potentially involved in P catabolism, based on our genomic and proteomic analyses (Figure 4C and S5). SusD1, SusD2 and to a lesser extent SusD3 are more highly conserved than SusD4/5/6, indicated by very short branch lengths in their respective clusters (Figure 4B). Concurrently, genes encoding the previously characterised Pi-insensitive SusCD complexes (Figure 4A) were typically found in operons with genes encoding various different stages of polysaccharide catabolism, for example, glycosyl hydrolases and carbohydrate-binding proteins found in the CAZyme database. Various predicted SusC/TBDT that did not have a corresponding SusD domain also responded to Pi-depletion. This included a TBDT associated with a 3-phytase (PusC8) in these *Flavobacterium* isolates as well as one whose gene is located adjacent to *phoBR* (PusC9, Fjoh_0545 in DSM2064) (Figure 4A). The genes corresponding to some PusC-like proteins were co-localised with predicted sugar phosphatase permease-like transporters which were also expressed during Pi depletion, as well as found in operons with highly expressed lipoproteins discussed next.

**Figure 4.**
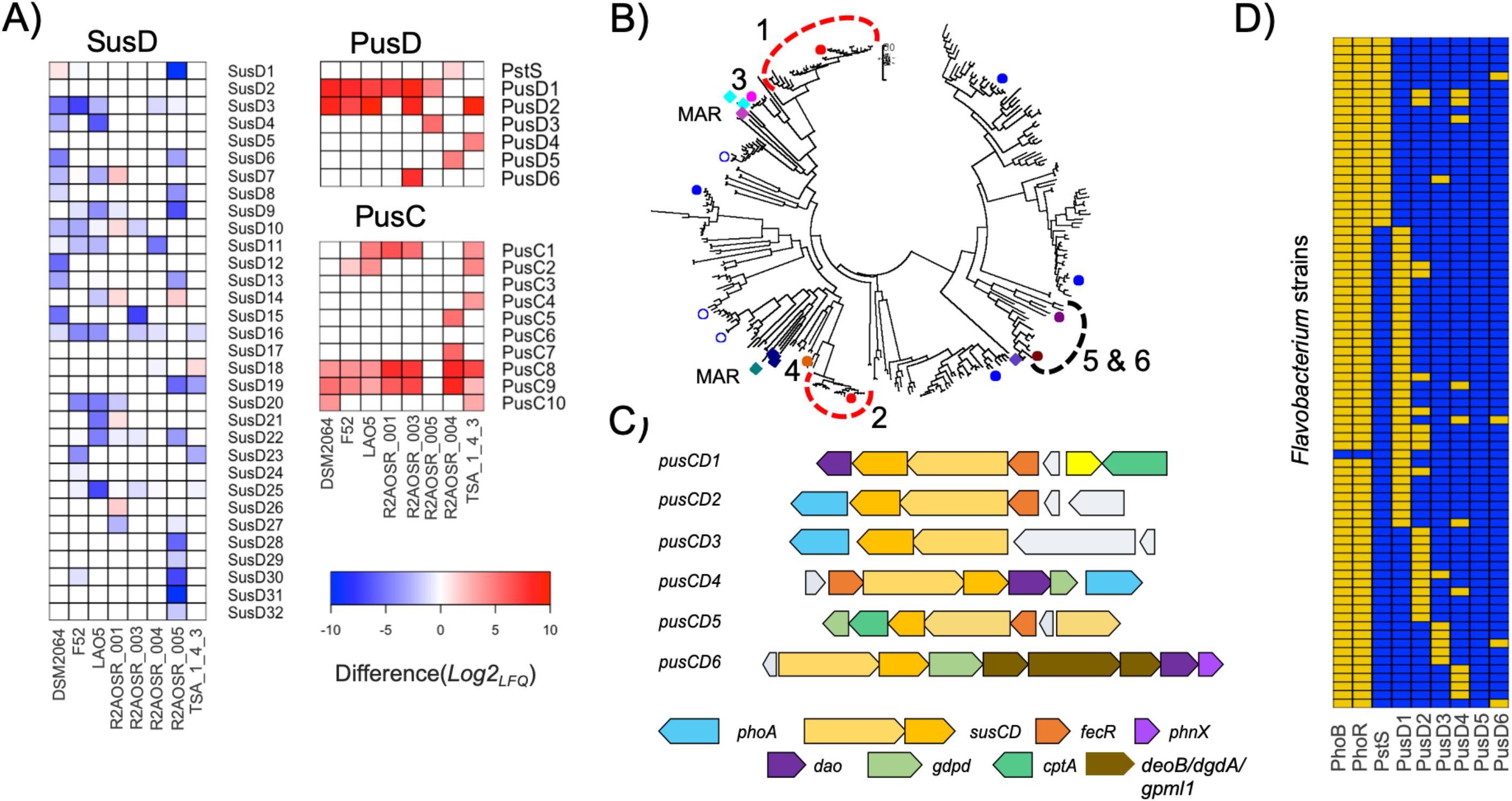
**(A)** The differential expression of SusCD complexes in the exoproteomes (n=3) of *Flavobacterium strains*, including Pi-insensitive (left column) and Pi-sensitive (right column). Red indicates an enrichment in Pi-deplete and blue indicates an enrichment in Pi-replete. For convenience, the difference in log2 LFQ intensities has been inversed compared to Figure 3, i.e. a positive difference equals an enrichment in Pi-deplete. **(B)** Maximum-likelihood tree displaying the phylogeny of various *Flavobacterium* SusD/PusD homologs. Marine (MAR) sequences identified in^32^ were added to the alignment and highlighted by squares. Certain non Pi-responsive *F. johnsoniae* DSM2064 SusD homologs are highlighted (blue circles). Red, maroon and pink circles represent the various Pi-sensitive PusD homologs. Numbers 1-6 represent the various PusD homologs. **(C)** The genetic neighbourhood of all PusCD complexes detected through exoproteomics, co-located with various genes predicted to be involved in Pi-acquisition. **(D)** Occurrence of various *pusD* and *pstS* genes in *Flavobacterium* genomes outlined in Table S12. Yellow squares indicate the presence of the given gene in a genome. Abbreviations: *phoA*, alkaline phosphatase A-like; *pusCD*, Pi-sensitive outer membrane SusCD-like complexes; *fecR*, iron-like transcriptional regulator; *phnX*, phosphonoacetaldehyde hydrolase; *dao*, oxidoreductase; *gdpd*, glycerolphosphodiesterase-like; ctpA, protease-like; *deoB*, Phosphopentomutase; *dgdA*, 2,2-dialkylglycine decarboxylase; *gpmI1*, 2,3-bisphosphoglycerate-independent phosphoglycerate mutase.

### *Flavobacterium strains* abundantly express three related hypothetical lipoproteins in response to Pi-depletion

During growth under Pi-deplete conditions, most *Flavobacterium* strains abundantly expressed three related hypothetical lipoproteins of unknown function, termed FI, FII and FIII. SWISS homology modelling showed that all three forms have structural features tentatively associated with pectate lyases/polygalacturonases (Figure 5A). However, phylogenetic analysis showed that these Pi-responsive forms are very different to characterised bacterial and fungal polygalacturonases and their apparent ‘true’ orthologs in *Flavobacterium* (Figure 5B). In DSM2064 (Fjoh_3856) and L0A5 (KNELKNMD_01352), FI was the most abundant protein detected in exoproteomes of Pi-deplete cultures. FI was expressed by all other isolates, albeit at a lower abundance, apart from F52 and WB32, which both lacked the corresponding gene. The gene encoding F1 was co-localised in a six-gene operon, which contained various predicted exoproteins, and was entirely expressed during Pi-depletion in the majority of strains (Figure 3, orange circles and squares and Figure 5C). FII (Fjoh_0546 in DSM2064) was a major constituent of the *Flavobacterium* exoproteomes during Pi-depletion. The gene encoding FII was co-localised with *phoBR*, *oprP* and another *tbdt* (also expressed during Pi depletion) (Figure 5D). In *F. johnsoniae* DSM2064, F52 and TSA_1_4_3, another gene was located at the end of this cluster and the corresponding hypothetical lipoprotein it encodes was also a major constituent of their exoproteomes during Pi-depletion (Figure 3, pink circles, and Figure 5D). The third form (FIII - Fjoh_4889 in DSM2064) was also expressed in the majority of strains during Pi depletion, but at a lower abundance.

**Figure 5.**
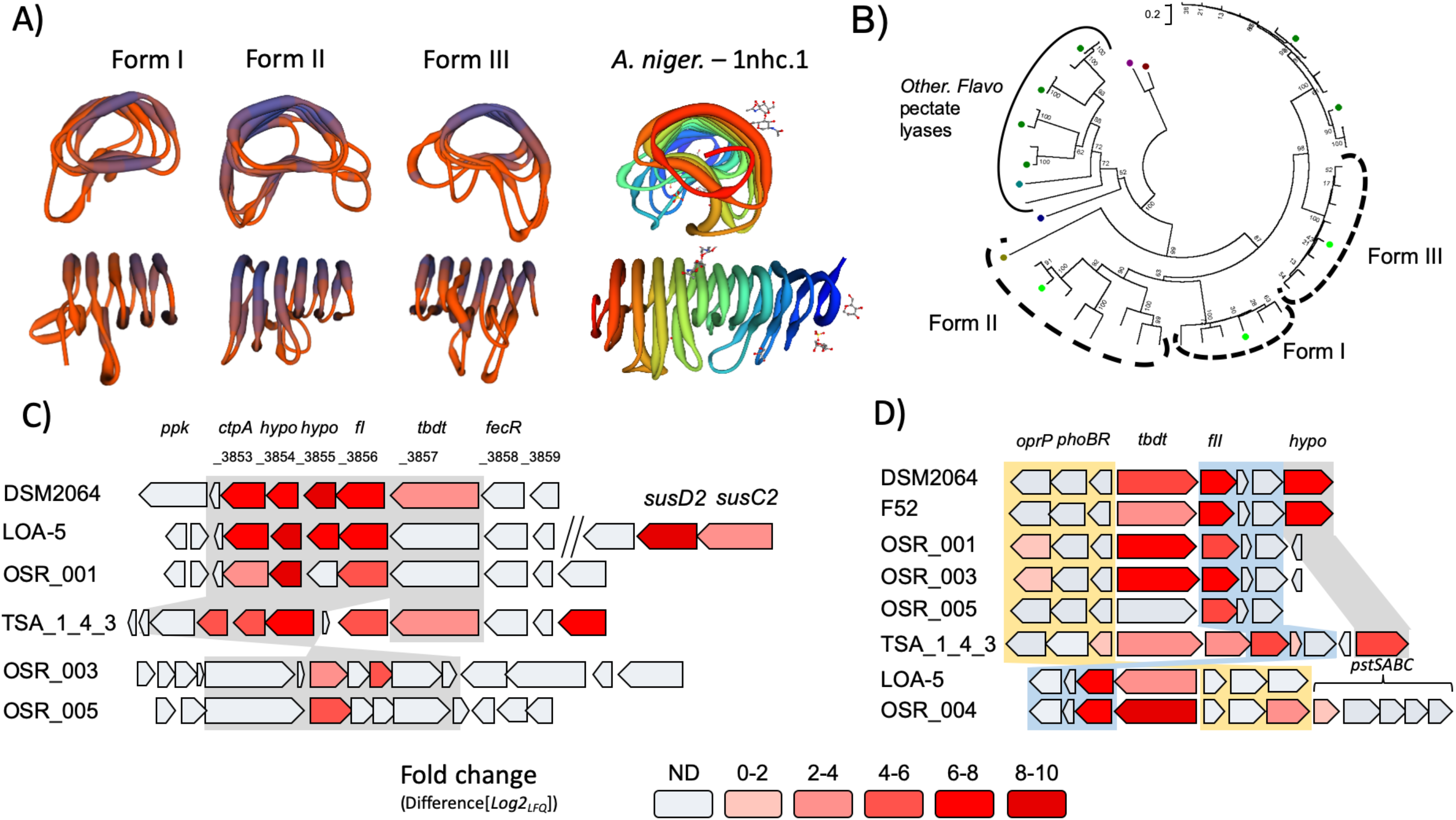
**A)** Structural modelling of the abundantly secreted Pi-sensitive hypothetical exoproteins, FI, FII and FIII. SWISS homology modelling revealed that these hypothetical exoproteins share structural homology with pectate lyases/ characterised polygalacturonases found in Bacteria and Fungi. **(B)** Maximum-likelihood tree displaying the phylogeny of low-Pi inducible hypothetical exoproteins FI, FII, and FIII (light green represents DSM2064). The previously characterised classical polygalacturonases (Maroon, Dark Blue and Mustard) and other predicted polygalacturonases found in DSM2064 and F52 (Dark green) were also added to the analysis. **(C)** The genetic neighbourhood of the gene encoding FI, which was the most abundant protein in Pi deplete exoproteomes of DSM2064 and LAO5, illustrating co-localisation with various other low Pi-responsive genes in a genetic operon. **(D)** The gene encoding FII was co-localised with genes encoding the PHO regulon two-component regulatory system for the PHO regulon, *phoBR*, a Pi-sensitive *tbdt/susC*-like transporter, a putative outer membrane phosphate selective *oprP*. For OSR_004, the genes encoding the high affinity transport system, *pstSABC* were also co-localised in this region. For convenience, the difference in log2 LFQ intensities has been inversed compared to Figure 3, i.e. a positive difference equals an enrichment in Pi-deplete.

**Figure 6.**
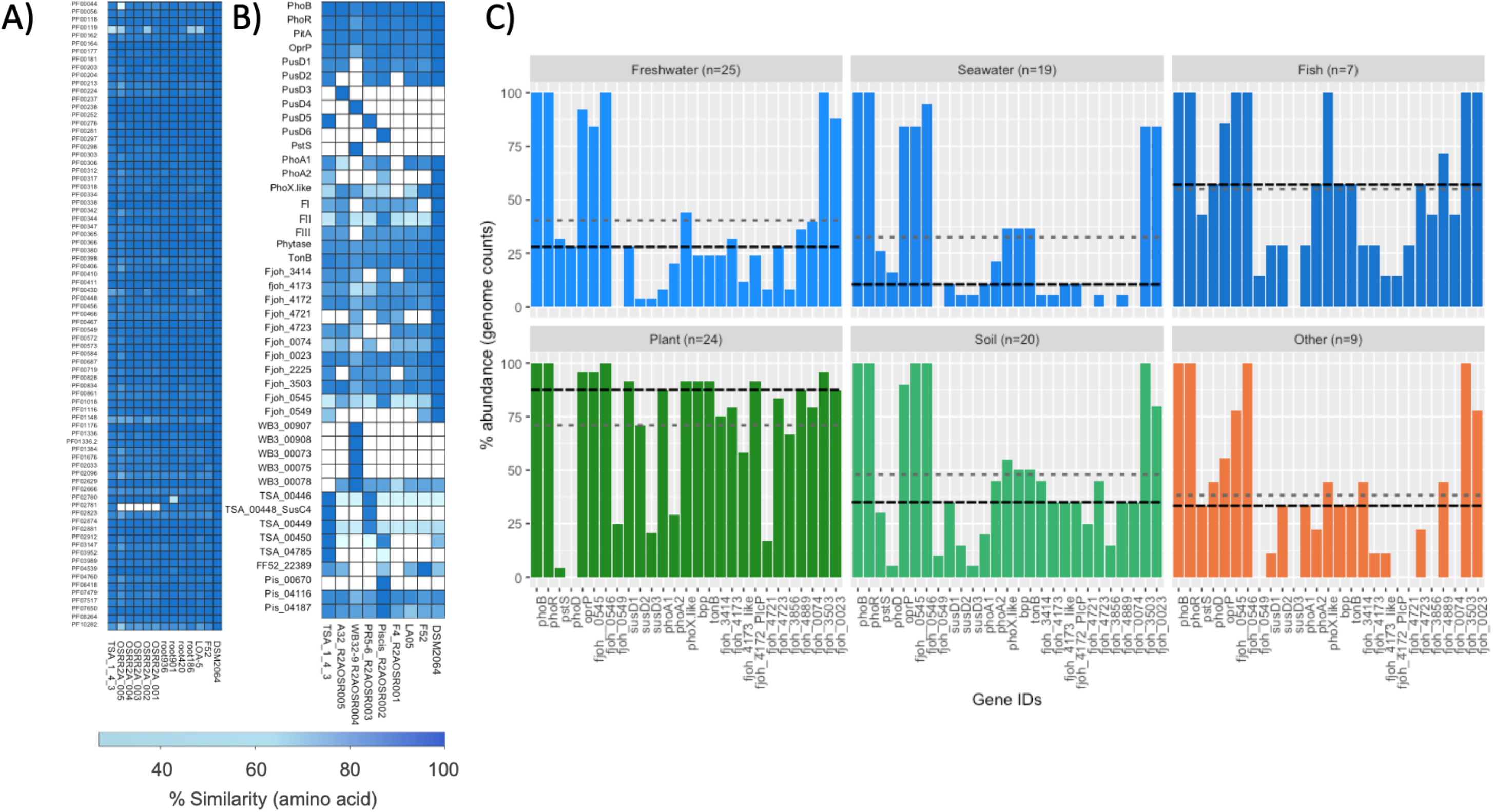
Genomic comparison of selected open reading frames (ORFs) amongst the *Flavobacterium* strains used in this study. **(A)** ORFs representing housekeeping genes and core metabolic genes identified by^76^ were used to determine genome completeness and isolate similarity. All genes were compared against DSM2064 queries. **(B)** Genes encoding all the potential P acquisition proteins as well as the master regulator PhoBR were compared. The scale bar represents the percentage identity at the amino acid level of each orthologue. White squares indicate no orthologue in a given genome. **(C)** The presence of PHO regulon genes in >100 *Flavobacterium* genomes obtained from different environments. *Flavobacterium* genomes were partitioned based on their source of isolation. n= the number of genomes analysed from each environmental niche. The mean (light grey dashed line) and median (black dashed line) values for the percentage abundance of all PHO regulon are shown for each graph.

Taken together, these exoproteomic data reveal there are numerous novel extracellular enzymes and transporters linked to the *Flavobacterium* PHO regulon. This demonstrates that this genus has unique mechanisms to acquire P similar to their distinct carbon acquisition strategies.

### Genomic comparison of the *Flavobacterium* PHO regulon among other *Bacteroidetes* isolates

Compared to their core metabolic and housekeeping genes, the eight *Flavobacterium* strains showed a high degree of variation in their P acquisition genes (Figure 6A and B). Whilst most strains possessed *susD1*, *oprP*, *bpp* (encoding a beta-propeller phytase), the majority of other genes (e.g *phoA1*, *phoA2*, *phoX*, *fjoh_4721/3, fjoh_0545*, *fII*) showed either an inconsistent distribution among the isolates or higher sequence variability (Figure 6B, full details in S13). OSR_004, which was the most phylogenetically distinct isolate, also showed least conservation of the PHO regulon. The variability in PHO regulon associated genes was not mirrored in the variability of housekeeping genes (*gyrB*, *recA*, *dnaK*) or *phoBR*, which all maintained a very high level of sequence similarity between the eight isolates.

To better understand the conservation of these low Pi-responsive genes/proteins throughout the *Bacteroidetes* phylum, we again analysed the 100 *Flavobacterium* genomes. We partitioned the 100 *Flavobacterium* genomes based on their broad environmental niche, e.g. freshwater, seawater, soil, fish and plant-associated, which included rhizosphere, phyllosphere and endosphere. Almost all of the genes encoding the low Pi-responsive proteins detected in our exoproteomic analysis were enriched in the genomes of plant-associated genomes compared to the other environmental niches (Figure 6C). Therefore, these data suggest that rhizosphere-dwelling *Flavobacterium* possess unique niche-associated genes for mobilising and acquiring P.

### *Flavobacterium* possess novel phosphatases resulting in molecular redundancy

Based on our exoproteomic data we hypothesised the PhoX-like protein (encoded by *fjoh_2478*, hereafter termed *phoX:2064*), abundantly expressed under Pi depletion in all strains, was responsible for the inducible APase activity during Pi depletion. However, mutagenesis of *phoX:2064* only reduced APase activity during Pi-deplete growth by 33% (Figure 7A). To determine which other APases were responsible for the residual activity *in vivo*, we heterologously expressed several other putative *Flavobacterium* APases, as well as *phoX:2064*, in an APase knockout mutant of *Pseudomonas putida* BIRD-1 (*phoX:BIRD-1*). These candidate *Flavobacterium* APases consisted of two genes (*fjoh_3187* and *fjoh_3249*) encoding the PhoA-like proteins, as well as the putative phosphatases encoded by *fjoh_3414* and *fjoh_4173*. The native *phoX:BIRD-1* and *phoA* from *E. coli* were used as controls. The native *phoX:BIRD-1* and *E. coli phoA* both restored APase activity in the *Pseudomonas* mutant (Figure 7B). However, neither *phoX:2064*, nor *fjoh_3414* and *fjoh_4173* produced any detectable APase activity. Both *phoA1* (*fjoh_3249*) and *phoA2* (*fjoh_3187*) restored APase activity in the *Pseudomonas* null mutant, demonstrating these forms are *bona fide* APases. Subsequent mutation of both *phoA* genes in *F. johnsoniae* resulted in a 36% reduction in Pi-deplete APase activity suggesting that they are expressed during Pi-depletion, despite the fact that PhoA1 was not detected in the exoproteome of *F. johnsoniae* DSM2064 and PhoA2 was at very low abundance (Table S3). Finally, we hypothesised that another lipoprotein, predicted to be a member of the endonuclease/exonuclease/phosphatase superfamily, encoded in *F. johnsoniae* DSM2064 by *fjoh_0074*, may play a role in the constitutive APase activity observed during Pi-replete as well as Pi-deplete conditions. A quadruple knockout mutant (*Δfjoh_0074:Δfjoh_2478: Δfjoh_3249: fjoh3187*) showed a 53% reduction in Pi deplete APase activity but no significant effect on Pi replete APase activity (Figure 7A). We confirmed residual APase activity in our various mutants, as well as our heterologous expression *Pseudomonas* strains, using a plate screening method, where a blue colour is indicative of APase activity (Figures S6 & S7). In summary, we identified three APases (PhoX-like, PhoA1, PhoA2) that are collectively responsible for approximately half of the observed Pi-deplete APase activity in *F. johnsoniae*, although the role of Fjoh_0074 is still uncertain. Thus, *F. johnsoniae* clearly expresses unique and, as yet unidentified APases that are unlike any previously characterised forms.

**Figure 7.**
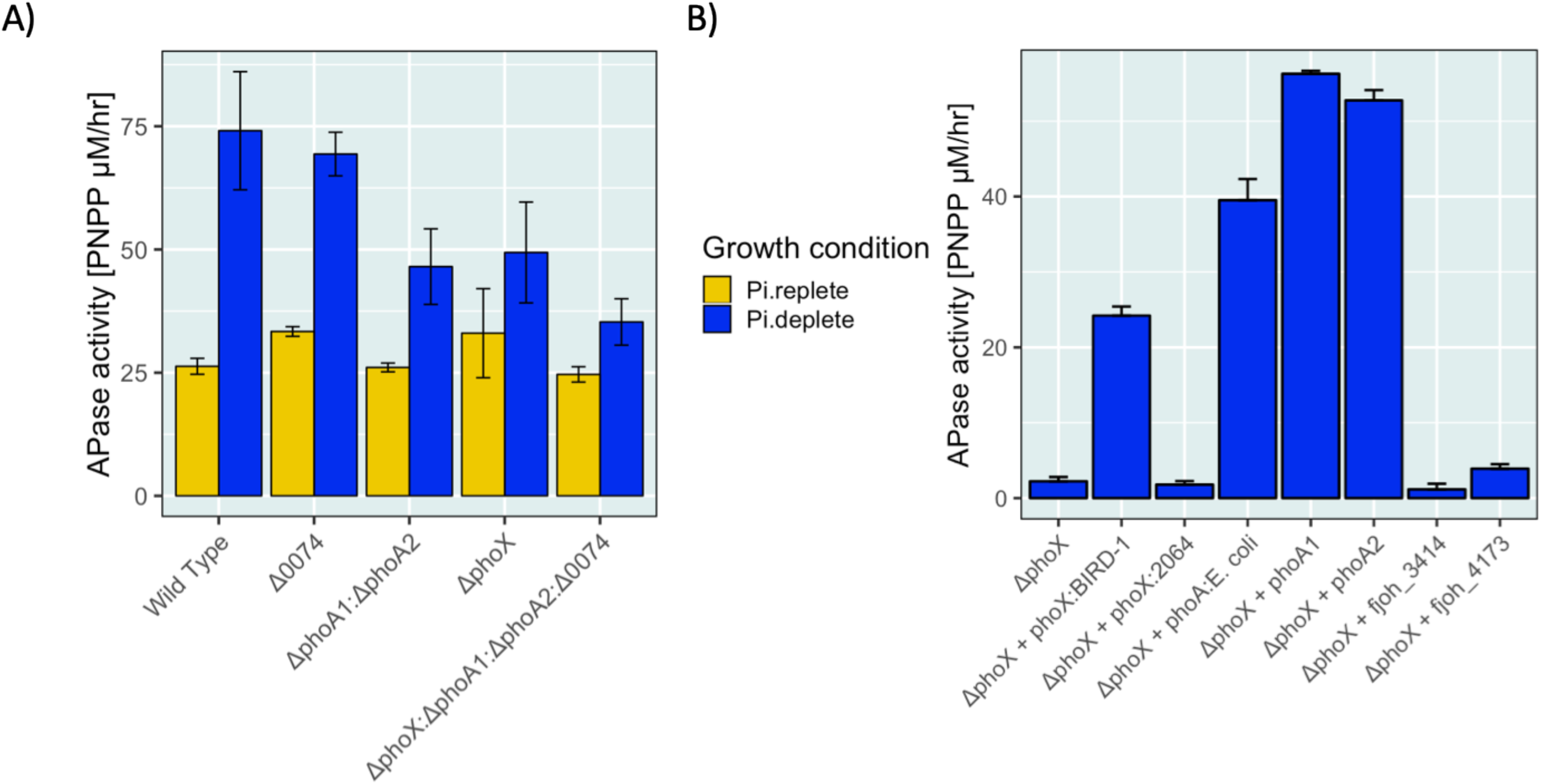
**(A)** Phosphatase (monoesterase) activity (n=3) of *F. johnsoniae* DSM2064 genetic variants, including the wild type during Pi-replete and Pi-deplete growth conditions. Para-nitrophenol phosphate (PNPP) was used as the substrate. **(B)** Complementation of the *ΔphoX Pseudomonas putida* BIRD-1 mutant with different candidate phosphatase genes from *F. johnsoniae* DSM2064. *phoX* from *Pseudomonas* sp. BIRD-1 (PPUBIRD-1_1093) and *phoA* from *E. coli* were used as controls. Note that the *phoX*-like gene (Fjoh_2478) from *F. johnsoniae* DSM2064 did not restore APase activity in the mutant, whilst the two *phoA*-like (Fjoh_3249 and Fjoh_3187) genes did. Results presented are the mean of triplicate assays from three independent cultures. Error bars denote standard deviation.

### Low-Pi inducible SusCD complexes in *Bacteroidetes*: A novel mechanism for P uptake using SusCD in *Bacteroidetes*?

As we identified several SusCD complexes (PusCD) whose expression was induced during Pi-depletion along with other co-localised genes predicted to act as phosphoestereases, we hypothesised that these transporters play a role in P acquisition in a similar manner to PstSABC or other ATP-binding cassette (ABC) transporters such as PhnDEF (phosphonates) or UgbABC (phosphodiesters). We screened 100 phylogenetically distinct plant-associated bacterial genomes (Table S12) and discovered that all non-*Bacteroidetes* strains possessed *PstS*, whilst the various *pusD* forms were exclusive to *Bacteroidetes* genomes (Figure 8A). We therefore mutated *pusCD1* and *pusCD2* in *F. johnsoniae* to determine the role of these outer membrane complexes in Pi-acquisition. Mutation of both *pusCD* complexes (*ΔpusCD1:ΔpusCD2*) did not result in any statistically significant differences in growth rate across a range of Pi concentrations compared to wild type (Figure 8B). In contrast, direct growth competition experiments between the mutant and wild type showed a small, but consistent fitness defect of the *ΔpusCD1:ΔpusCD2* mutant compared to the *F. johnsoniae* DSM2064 wild type across a range of Pi concentrations strongly implicating these PusCD complexes play a role in Pi acquisition.

**Figure 8.**
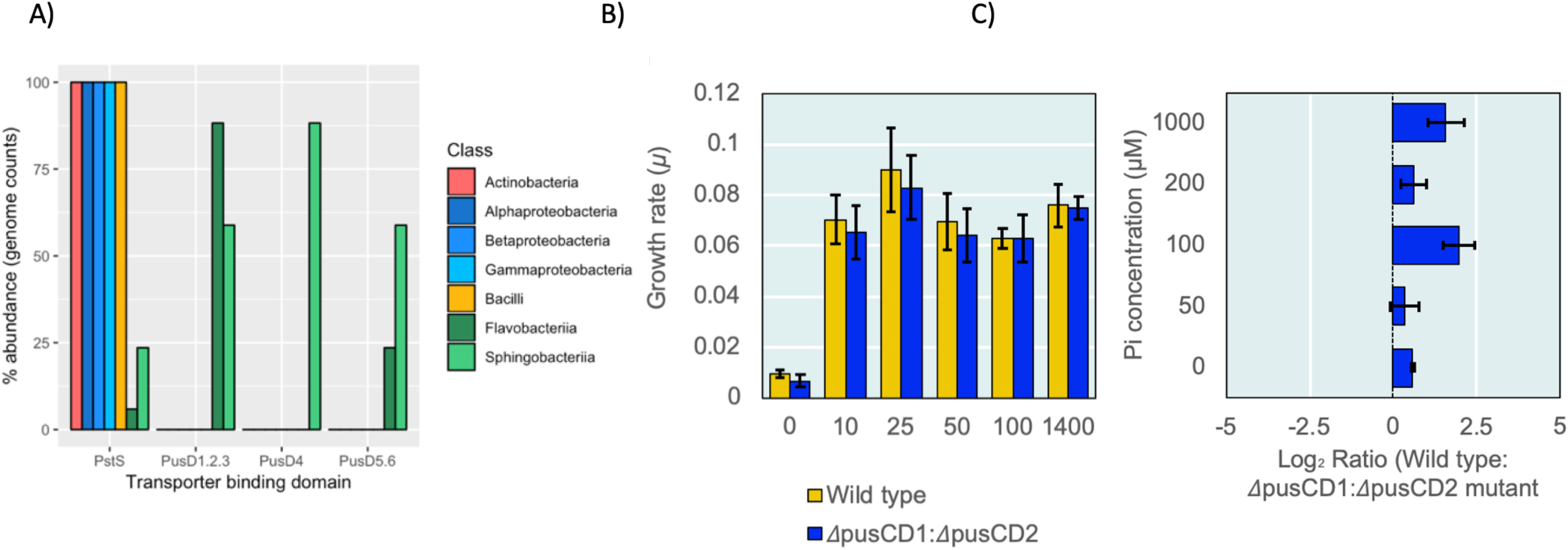
Characterisation of the (*ΔpusCD1:ΔpusCD2*) *F. johnsoniae* DSM2064 mutant. **(A)** Distribution of either genes encoding PstS or the various PusD homologs in 100 rhizobacterial genomes (Table S11) chosen from IMG/JGI. Genomes were categorised according to class to differentiate between *Flavobacteria* and *Sphinogbacteria* **(B)** Growth rates of either the mutant or wild type grown on glucose (20 mM) and various concentrations of Pi. **(C)** Competition experiments (n=3) between the mutant and wild type. Results plotted are the log2 of the ratio between wild type and mutant colony forming units (c.f.u.). Final cell counts varied from 106 to 109 c.f.u across the Pi concentrations gradients. Error bars denote standard deviation.

## Discussion

*Flavobacteriaceae* have recently been shown to be major constituents of the plant microbiome yet we know very little about their physiological and thus functional role within this niche^20,28,48^. Whilst recent evidence has demonstrated they have a potentially unique role in rhizosphere carbon cycling^27^, knowledge regarding their potential to mobilise P is very limited. Therefore, our ability to determine their importance and potential for exploitation remains unclear. Here, we observed that *Flavobacterium* isolates exhibit superior APase activity compared to *Pseudomonas*, especially at lower pH ranges more aligned to conditions in the rhizosphere (Figure 1A). Moreover, we also discovered that *Flavobacterium* exhibited APase activity irrespective of excess Pi concentrations (Fig. 1A), which is a unique and potentially important trait in the context of sustainable agriculture^6,7,40^.

We therefore sought to determine the components of the *Flavobacterium* PHO regulon, one of the cell’s major regulatory networks that helps to mobilise different forms of recalcitrant P_o_ and P_i_ ^8^. Utilising exoproteomics^49^, we revealed that *Flavobacterium* isolates express novel genes and gene clusters which we predict may access complex forms of P_o_ within the rhizosphere. In support of this, the *Flavobacterium* strains abundantly secreted numerous proteins whose genes were enriched in genomes of plant-associated isolates (Figure 6). In particular, the hypothetical lipoproteins FI (and associated gene cluster), FII and FIII, which were abundantly expressed in the majority of strains tested during Pi-depletion, are present in a significantly higher number of plant-associated *Flavobacterium* genomes compared to soil and aquatic strains, especially seawater (Figure 6B). In human and ocean microbiomes, *Flavobacteriaceae* and other member of the *Bacteroidetes* phylum are the dominant players in the degradation of recalcitrant plant or algal derived polymers ^23,25,26,31,50^. Given the fact FI, FII and FIII share structural homology with fungal and bacterial pectin lyases/polygalactouranases and are located in genetic clusters containing predicted Pi-related genes, our data suggests that these hypothetical lipoproteins maybe involved in the binding and subsequent degradation of complex plant-derived P-containing compounds. This hypothesis is further supported by the observation that several SusCD-like complexes, which are known to transport oligosaccharides and small polysaccharides^29–31,51,52^, are expressed during Pi-depletion.

To date, SusCD complexes have traditionally been shown to be involved exclusively in plant-derived polysaccharide transport and degradation^29,30,52^. Despite the fact that members of the *Bacteroidetes* phylum can possess over 100 *susCD* clusters in their genomes^31,53^, their functional characterisation has been slow. For the first time, we have identified specific SusCD complexes, termed here PusCD, that are expressed during Pi-depletion in a conserved manner across different *Flavobacterium* isolates. This suggests a novel role for these poorly characterised transport systems. Based on our exoproteomic, genomic and experimental data, we have two unresolved hypotheses for these transport systems: PusCD complexes are involved in 1) high affinity uptake of Pi, akin to the traditional PstSABC transport system 2) they have evolved to help plant-associated *Flavobacteria* acquire P from P_o_ complexes typically found in the rhizosphere. Our experimental data suggests that PusCD1/PusCD2 indeed play a role in Pi acquisition given the fitness defect of the *pusCD1*:*pusCD2* double mutant in competition experiments with the wild type across a range of Pi concentrations (Fig. 8C). Moreover, our exoproteomic and genomic data clearly showed that PusCD1 and PusCD2 were abundantly expressed in plant-associated *Flavobacterium* strains that lack *pstSABC*, which is the most conserved constituent of the bacterial PHO regulon across all taxa^8^ and the major high affinity Pi transporter in most bacteria e.g. *Pseudomonas^13,44,54^, Synechocystis*^55^, *Prochlorococcus*^56^, *Streptomyces^57^, Bacillus*^14^ and *Pelagibacterales*^9^. Phylogenetic analyses also revealed that SusD1 and SusD2 may have evolved to transport Pi as they share a high degree of within genus conservation, which is usually associated with a specialised role in binding ligands that are structurally simple^32^. A role for the Pi-induced TBDT/SusC-like proteins, lacking SusD partners, may explain the non-essential role of SusCD1 and SusCD2 in Pi acquisition. With respect to the second hypothesis, it is already known that human gut and marine-dwelling *Bacteroidetes* utilise SusCD complexes to transport larger oligosaccharides into the periplasm after initial cleavage of the polysaccharide at the outer membrane, resulting in little or no loss of the high molecular weight substrate to the environment^51,58^. Furthermore, in *Bacteroides thetaiotaomicron* Bt-8736, transport of the entire polysaccharide inulin was conferred by the presence of a distinct *susCD* complex located within a PUL, precluding the need for any extracellular degradation of the compound^52^. Given the fact that all the *Flavobacterium pusCD* complexes, particularly *pusCD4/5/6* are co-localised in operons with various phosphonatases, monoesterases and diesterases containing predicted signal peptide domains (Figure S5), a similar mechanism maybe operational for the degradation of complex P_o_ compounds. Degradation of complex P_o_ compounds for growth would be akin to their unique carbon acquisition strategies^23,26,27,59^ and may represent another example of niche partitioning in this phylum. Either hypothesis would be a completely novel mechanism for Pi transport, where ligand binding and active transport takes place at the outer membrane (as opposed to the periplasm and inner membrane using *pstSABC*), potentially concentrating the periplasm with Pi, that is subsequently transported to the cytoplasm through a permease, PitA.

P_o_ often accounts for the majority of total P in soils and in agricultural systems this is set to rise with the increased use of organic P fertilisers, primarily from manure and other waste materials^4^. Despite this, research and practices tend to focus on the availability of Pi and often neglect the P_o_ cycle^6,7^. PhoD and PhoX are frequently used as molecular markers to determine APase activity in agricultural soils^43,60,61^ due to their high expression and contribution to APase activity in traditional environmental laboratory isolates such as *Pseudomonas* ^13,44^, *Bacillus* ^62^ and *Sinorhizobium^63^*. For the first time, we have shown that rhizosphere-dwelling isolates related to *Flavobacterium* and *Bacteroidetes* more generally express constitutive APase activity even in the presence of excess Pi. Therefore, *Bacteroidetes* offer potential novel solutions to aid in the recovery of Pi in soils and wastewater treatment plants. Interestingly, although the abundantly secreted *Flavobacterium* PhoX, which is phylogenetically distinct from previous characterised forms (Figure S2), was responsible for part of the APase activity under Pi-depletion, this protein did not restore APase activity in a *Pseudomonas* APase null mutant (Figure 7B). Unlike, *Pseudomonas* PhoX, which is translocated to the outer membrane through interaction of the Twin-Arginine Translocation (TAT) pathway and type II secretion pathway^64^, *Flavobacterium* PhoX contains a signal peptide associated with secretion via the sec pathway. However, how *Flavobacterium* PhoX is transported to the outer membrane remains unknown. *Flavobacterium* PhoX homologs do not contain the C-terminal domains (CTD) associated with secretion via the type IV secretion system found in numerous Bacteroidetes including *F. johnsoniae^47,65^*. Understanding, the secretion mechanism of PhoX and other secretory APases is essential for exploring their potential discovery in functional metagenomics screening approaches. Interestingly, only the two PhoA-like homologs elicited heterologous APase activity in the *P. putida ΔphoX* mutant despite the fact that their expression was patchy and weak across the eight isolates tested. PhoA is generally considered to be associated with enteric bacteria and other human-associated bacteria^16,66^. Thus, it is interesting that *phoA1* and *phoA2* are enriched in plant-associated genomes. The fact that our quadruple APase mutant only resulted in a ~50% reduction in inducible APase activity (Figure 7A), suggests *Flavobacterium* possesses redundant and unique APases which may act on different fractions of P_o_ found in soil and the rhizosphere. From our exoproteomic dataset, there are at least 20-30 potential candidates that may explain the APase activity in *F. johnsoniae* DSM2064 which will require a significant amount of further investigation. It is tempting to speculate that under Pi-replete conditions *Flavobacterium* strains express APases to access the carbon associated with P_o_-containing compounds, which would ultimately mobilise Pi in the rhizosphere.

In conclusion, a combination of whole cell physiology, exoproteomics, comparative genomics and bacterial genetics revealed that *Flavobacterium*, which exhibits strong and constitutive APase activity, possesses unique molecular mechanisms for Pi-acquisition. In addition to expressing novel gene clusters with potentially unique substrates specificities, *Flavobacterium* likely express multiple uncharacterised APases. Furthermore, we have identified for the first time, a group of SusCD complexes that appear to play a role in Pi-acquisition. This study paves the way for further exciting research on the physiology of *Flavobacterium* strains and their potential exploitation as microbial tools in sustainable agriculture.

## Materials and Methods

### Isolation and growth medium used for cultivating Bacteroidetes

*Flavobacterium johnsoniae* DSM2064 (UW101) was purchased from the Deutsche Sammlung von Mikroorganismen und Zellkulturen (DSMZ). *Flavobacterium* sp. F52, which was isolated from the rhizosphere of bell pepper ^67^ and *Flavobacterium sp.* L0A5, which was isolated from the rhizosphere of a desert plant (Cytryn, unpublished data) were both kindly donated from the Cytryn Lab, Agricultural Research Organisation, Israel. Strains were routinely maintained on casitone yeast extract medium (CYE)^68^ containing casitone (3 g ^L-1^), yeast extract (1 g ^L-1^), MgCl_2_ (350 mg ^L-1^) and 30 g ^L-1^ agar. To isolate all other strains a modified R2A medium developed by^42^ was used. Briefly, the medium was made in two parts and autoclaved separately. Each solution comprised: Solution A, yeast extract (1.0 g L^−1^), tryptone (0.5 g L ^−1^), peptone (1.5 g L ^−1^), starch (1.0 g L ^−1^), glucose (1.0 g L ^−1^), and agar (30 g L ^−1^); Solution B, K_2_HPO_4_ (0.6 g L ^−1^), MgSO_4_ (48 mg L ^−1^), and sodium pyruvate (0.6 g L ^−1^). Cyclohexamide (100 mg L ^−1^) and Kanamycin (40 mg L ^−1^) were added to reduce fungal and non-Flavobacterial growth. To investigate the effect of Pi-depletion on the *Flavobacterium* strains, each was grown (n = 3) in a Minimal A medium adapted from^13^. Briefly, the medium contained: Glucose 3.6 g ^L−1^, NaCl 200 mg ^L−1^, NH_4_Cl 450 mg ^L−1^, CaCl_2_ 200 mg ^L−1^, KCL mg ^L−1^ MgCl_2_ 450 mg ^L−1^, FeCl_2_ 10 mg ^L−1^, MnCl_2_ 10 mg ^L−1^, 20mM Bis/Tris buffer pH 7.2, KH_2_PO_4_ added to a final concentration of either 50 μM or 1.4 mM. Each strain was pre-cultured in minimal A medium containing 400 μM Pi to ensure cells had adequate Pi whilst minimising the potential for residual Pi carry over into the triplicate experimental cultures.

### Isolation of Bacteroidetes

Four isolation, two fields trials growing *Brassica napus* L. (OSR) were utilised, the first was at located Warwick HRI (April, 2017), University of Warwick and the other was located at Sonning Farm, University of Reading (Sept, 2017). Samples were collected during flowering stage at Warwick HRI and during the four-leaf stage at Sonning Farm. Plants were removed from soil and shaken to remove all loosely adhering soil. Roots containing tightly adhering soil were cut and washed (shaken vigorously) in 10 mL phosphate buffer saline solution to remove rhizosphere soil. A serial dilution of the resultant soil solution was set up and plated on to either modified R2A or CYE. Individual colonies were picked and sub cultured prior to identification by 16S rRNA gene analysis.

### Competition experiments between *F. johnsoniae* DSM2064 wild type and the *susCD1/2* mutant

Each strain was grown overnight in CYE. Cells were pelleted and resuspended in the same volume of the defined minimal A medium containing no Pi. For each strain, the cell suspension was used to inoculate (1 % v/v) flasks (n=3), in a 1:1 ratio. containing either no Pi, 50 μM, 100 μM, 200 μM or 1 mM Pi. After overnight growth, cultures were sub-cultured using 2% (v/v) of the same medium. Colony forming units were calculated by plating serial dilutions of the culture onto LB. To distinguish between the wild type and mutant, we took advantage of a previous wild type variant that had lost the majority of its yellow pigmentation, but had no observable effect on growth. Therefore, we constructed the susCD knockouts in this strain variant. Plates counts were performed for T0 to ensure the correct 1:1 ratio was achieved.

### Quantification of alkaline phosphatase activity

The protocol was identical to Lidbury et al., (2016). Briefly, a 0.5 mL culture (n = 3) was incubated with 20 μL *para*-nitrophenyl phosphate (pNPP) (final conc. 4 mM) and incubated at room temperature until colour development started to occur (roughly 5-20 min). The reaction was stopped using 25 μL 2 mM NaOH and incubated for 10 min. Cell debris and precipitants were removed via centrifugation (2 min, 8,000 × g) prior to spectrophotometry (405 nm). A standard curve for para-nitrophenol was generated using a range of known concentrations (0, 4, 8, 25, 50, 75, 100 mg mL^−1^).

### Exoproteomic analysis of the *Flavobacterium* isolates

For each isolate, 50 mL cultures (n=3) were grown overnight (~16 h) under either Pi-deplete (50 μM) or Pi-replete (1.4 mM) conditions. To capture the exoproteome we followed the method described by Lidbury et al., (2016)^13^. Briefly, cells were pelleted at 4,000 rpm @ 4°C in a Beckman benchtop centrifuge prior to gentle hand filtration through 0.45 μm and 0.22 μm PDVF membranes. The remaining supernatant was frozen at −20°C until further analysis. Culture supernatants were thawed over night at 4°C and exoproteins were concentrated following the Trichloroacetic acid precipitation protocol described by Christie-Oleza et al., (2012)^69^, with a slight modification. Protein pellets were dissolved in 60 μL Sample Buffer (Expedeon, UK) containing the RunBlue^©^ DTT Reducer (Expedeon, UK) following the manufacturer’s guidelines. Exoproteomes were visualised on a 1D-SDS PAGE gel (Expedeon, UK) using the manufacturer’s recommended settings. For protein identification a short run (~2 min) was performed to create a single gel band containing the entire exoproteome, as previously described by Christie-Oleza et al., (2012)^59^. In-gel reduction was performed prior to trypsin digest as previously described by^69^. Tryptic peptide mixtures were cleaned up according to the protocol of Kaur et. Al (2018)^70^. Samples were analysed by means of nanoLC-ESI-MS/MS using an Ultimate 3000 LC system (Dionex-LC Packings) coupled to an Orbitrap Fusion mass spectrometer (Thermo Scientific, USA) using a 60 min LC separation on a 25 cm column and settings as previously described^71^. Quantification, statistical analyses and data visualisation of exoproteomes was carried out in Perseus^72^. The mass spectrometry proteomics data have been deposited to the ProteomeXchange Consortium via the PRoteomics IDEntifications (PRIDE) partner repository with the dataset identifier PXD014380 and 10.6019/PXD014380.

### Genome sequencing of the *Flavobacterium* isolates

The OSR isolates were grown on CYE and cells were collected from plates by scraping of bacterial colonies. Cells were then sent to the Microbes NG unit (https://microbesng.uk/) at the University of Birmingham. All sequence data is deposited in the European Nucleotide Archive (ENA) database. For assistance, amino acid sequences for the complete CDS profile of all newly isolated strains are attached to the supplementary tables file. Raw contigs are also available on request.

### Mutagenesis of *Flavobacterium johnsoniae* DSM2064

To construct the various phosphatase and *pusCD* mutants in DSM2064, the method developed by^73^ was used. A full list of primers used in this study can be found in Table S14. Briefly, two 1-1.6 kb regions flanking each gene/gene operon/gene pair were cloned into plasmid pYT313^74^ using the HiFi assembly Kit (New England Biosciences). Sequence integrity was checked via sequencing. The resultant plasmids were transformed into the donor strain *Escherichia coli λ*S17-1 and mobilised into *Flavobacterium* via conjugation (overnight at 30°C). The donor: recipient suspension (1:1), was plated onto CYE containing erythromycin (100 μg mL^−1^). Kanamycin (50 μg mL^−1^) was used to counter selection against the donor strain. Single homologous recombination events were confirmed by PCR prior to overnight growth in CYE followed by plating onto CYE containing 10% (w/v) sucrose to select for a second recombination event. To identify a mutant, colonies were re-grown on CYE containing 10% (w/v) sucrose and CYE containing erythromycin 100 μg mL^−1^. Erythromycin sensitive colonies were screened for a double homologous recombination event by PCR targeting a region deleted in the mutant.

### Complementation of the *Pseudomonas ΔphoX* mutant

To complement the *Pseudomonas* sp. BIRD1 *ΔphoX* mutant with various APases, the promoter for the native *phoX*:BIRD-1, was cloned into the broad-host range plasmid pBBR1MCS-km^75^. The various APases were then subsequently cloned downstream of this promoter via Gibson cloning. All plasmids were mobilised into the *ΔphoX* mutant via electroporation using a voltage of 18 kv cm^−1^. Cells were immediately added to LB and incubated for 2-3 h prior to selection on LB supplemented with 50 μg mL^−1^ Kanamycin. All subsequent experiments were performed using LB containing 50 μg mL^−1^ Kanamycin to ensure strains kept the plasmids.

### Bioinformatics analysis

To determine the abundance of *phoA*, *phoD*, *pstS, phoX* genes in our newly isolated *Flavobacterium* strains, traditional BLASTP searches were performed using previously characterised genes. In some instances, certain genes were too distinct between different taxonomic groups, so HMMer was used to perform a hmmsearch based on profile Hidden Markov Models (pHMMs) (see Lidbury et al., 2017). In addition, a ‘function search’ was performed in IMG/JGI on either DSM2064 or *Flavobacterium* sp. F52 whose genomes have been previously uploaded to this database. The pfam domain search option was used to identify if genes contained predicted pfam domains, such as Pfam05787 for PhoX and Pfam00249 for PhoA. If DSM2064 or F52 had these orthologs these were used as query sequences to BLASTP against the OSR isolates and LAO-5. Given the fact these isolates all belong to the same genus, a high stringency was set (E^−90^) to try and remove any false positive predictions. Depending on the specific gene/protein these various methods were also used to determine the abundance of both previously characterised PHO regulon and newly identified low Pi-responsive genes/proteins in either the 100 *Flavobacterium* genomes or the 100 plant-associated taxa genomes. In the case of several genes, the predicted amino acid sequences were used to generate a phylogenetic tree to further determine the presence or absence of an ortholog as opposed to a paralog.

To determine genome completeness, orthologs of core and metabolic genes identified by^76^ were identified in DSM2064 and F52 using IMG/JGI. Local BLASTP was then performed on the OSR isolates and LAO-5 using DSM2064 orthologs as the query.

To determine potential protein structures of FI, FII and FIII, the online server interface for SWISS-model homolog analysis was utilised (https://swissmodel.expasy.org/). Default parameters were used to first determine suitable templates to build the model.

Phylogenetic analyses were performed using the software package MEGA7^77^. Alignment of the various protein sequences was achieved using MUSCLE. Phylogenetic distances were inferred using the p-distance model and evolutionary histories were inferred using the maximum-likelihood method based on the JTT matrix-based model.

## Acknowledgments

We would like to thank the Warwick Proteomics Research Facility, namely Dr. Cleidiane Zampronio and Dr. Juan Hernandez Fernaud for their assistance in generating and processing the mass-spectrometry data. We would also like to thank Prof. Mark McBride and Dr. Eddie Cytryn for kindly sending us various strains and plasmids used in this study. This study was funded by the Biotechnology and Biological Sciences Research Council (BBSRC) under the project code BB/L026074/1 linked to the The Soil and Rhizosphere Interactions for Sustainable Agri-ecosystems (SARISA) programme.

