## Supplementary materials 1 for "Novel mechanisms for phosphate acquisition in abundant rhizosphere-dwelling *Bacteroidetes*"

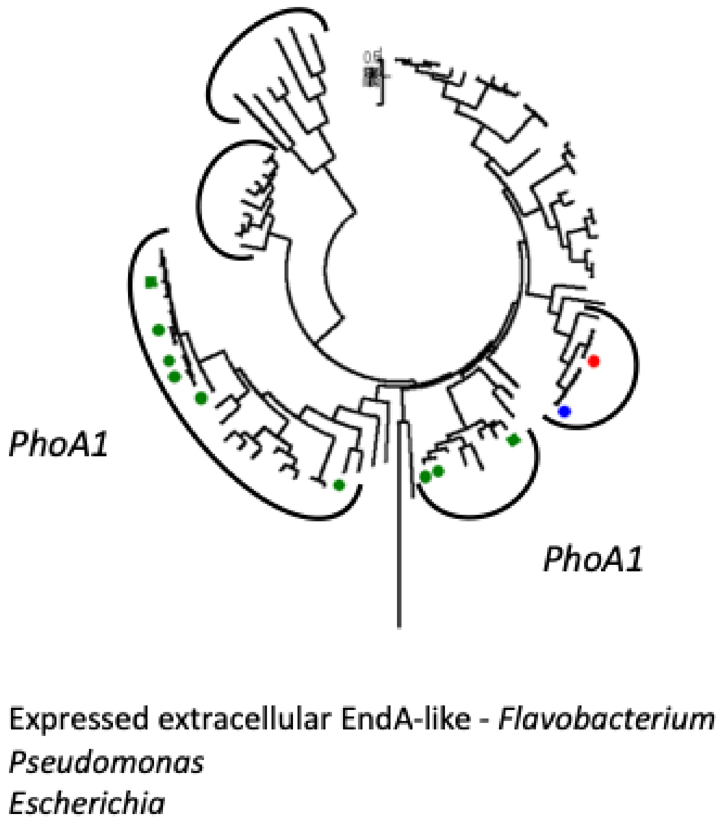


Figure S1. Maximum likelihood tree displaying the diversity of PhoA-like sequences. The evolutionary history was inferred by using the Maximum Likelihood method based on the JTT matrix-based model [1]. The tree with the highest log likelihood (-24607.22) is shown. Initial tree(s) for the heuristic search were obtained automatically by applying Neighbour-Join and BioNJ algorithms to a matrix of pairwise distances estimated using a JTT model, and then selecting the topology with superior log likelihood value. The tree is drawn to scale, with branch lengths measured in the number of substitutions per site. The analysis involved 90 amino acid sequences. All positions with less than 75% site coverage were eliminated. That is, fewer than 25% alignment gaps, missing data, and ambiguous bases were allowed at any position. There were a total of 262 positions in the final dataset. Evolutionary analyses were conducted in MEGA7 [2].

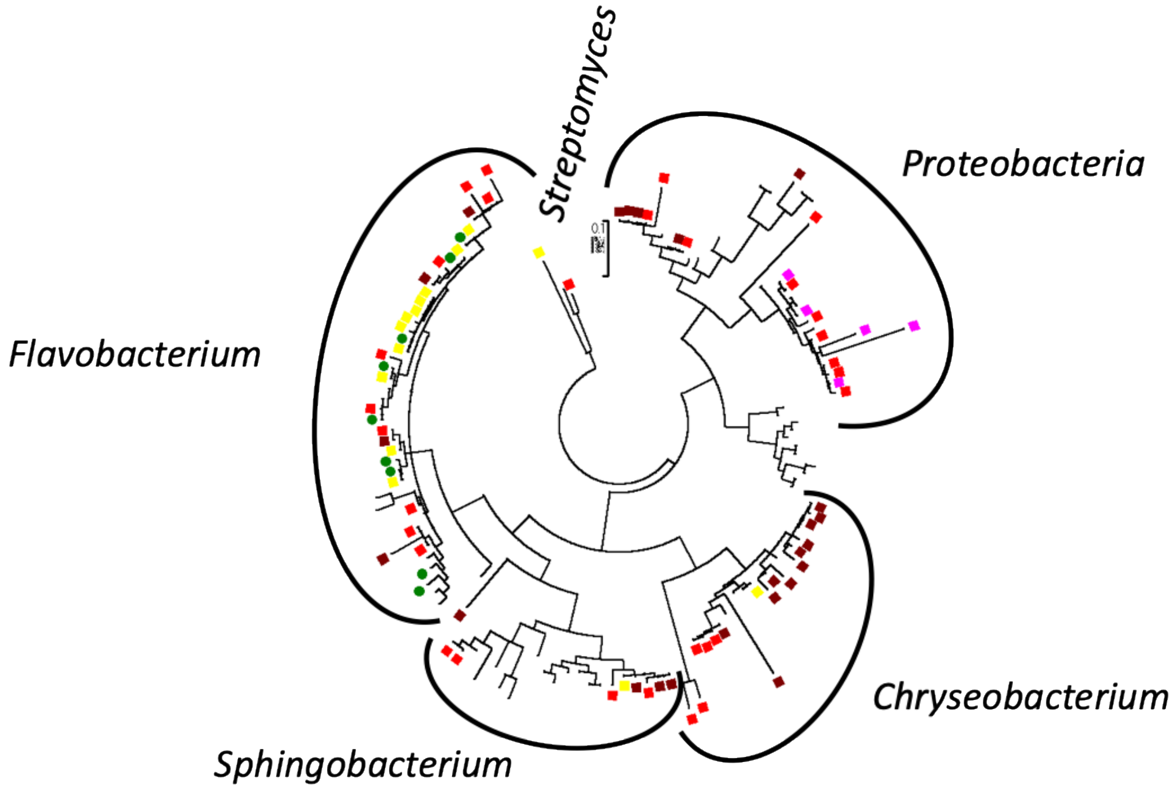


Figure S2. Taxonomy of the field-grown *Brassica napus* was determined using the 16S rRNA gene marker. The evolutionary history was inferred by using the Maximum Likelihood method based on the Tamura-Nei model [1]. The tree with the highest log likelihood (-10495.28) is shown. Initial tree(s) for the heuristic search were obtained automatically by applying Neighbour-Join and BioNJ algorithms to a matrix of pairwise distances estimated using the Maximum Composite Likelihood (MCL) approach, and then selecting the topology with superior log likelihood value. The tree is drawn to scale, with branch lengths measured in the number of substitutions per site. The analysis involved 134 nucleotide sequences. Codon positions included were 1st+2nd+3rd+Noncoding. All positions with less than 75% site coverage were eliminated. That is, fewer than 25% alignment gaps, missing data, and ambiguous bases were allowed at any position. There were a total of 433 positions in the final dataset. Evolutionary analyses were conducted in MEGA7 [2].

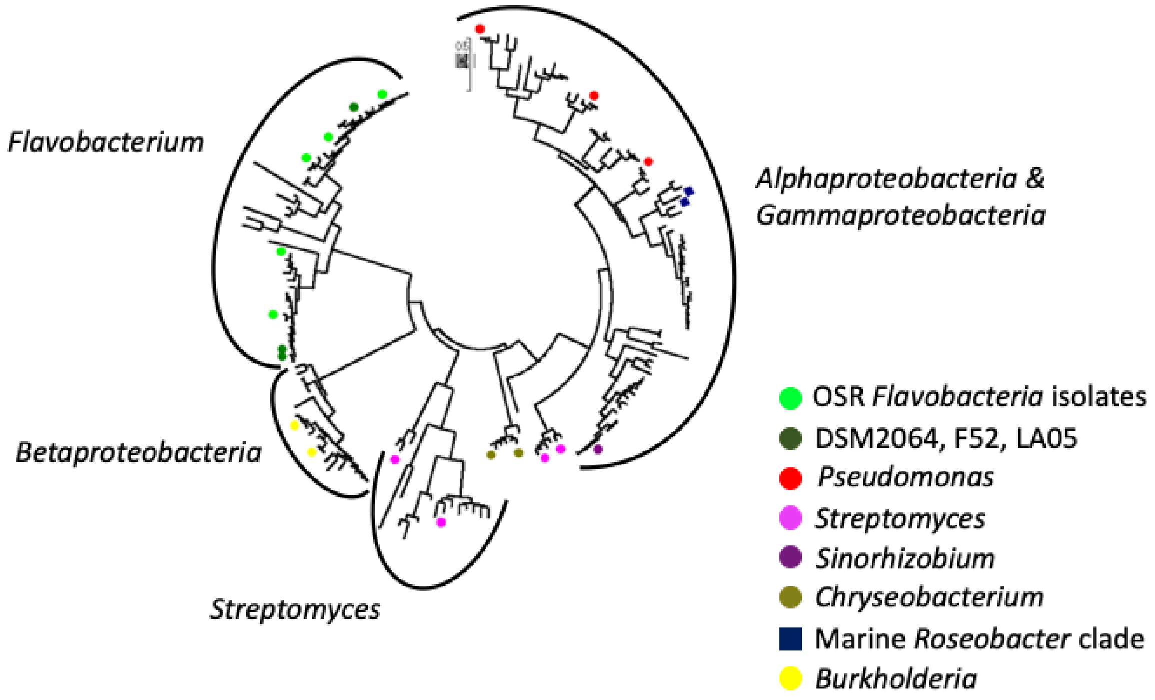


Figure S3. Maximum likelihood tree displaying the diversity of PhoX-like sequences. The evolutionary history was inferred by using the Maximum Likelihood method based on the JTT matrix-based model [1]. The tree with the highest log likelihood (-76880.03) is shown. Initial tree(s) for the heuristic search were obtained automatically by applying Neighbour-Join and BioNJ algorithms to a matrix of pairwise distances estimated using a JTT model, and then selecting the topology with superior log likelihood value. The tree is drawn to scale, with branch lengths measured in the number of substitutions per site. The analysis involved 302 amino acid sequences. All positions with less than 75% site coverage were eliminated. That is, fewer than 25% alignment gaps, missing data, and ambiguous bases were allowed at any position. There were a total of 277 positions in the final dataset. Evolutionary analyses were conducted in MEGA7 [2].

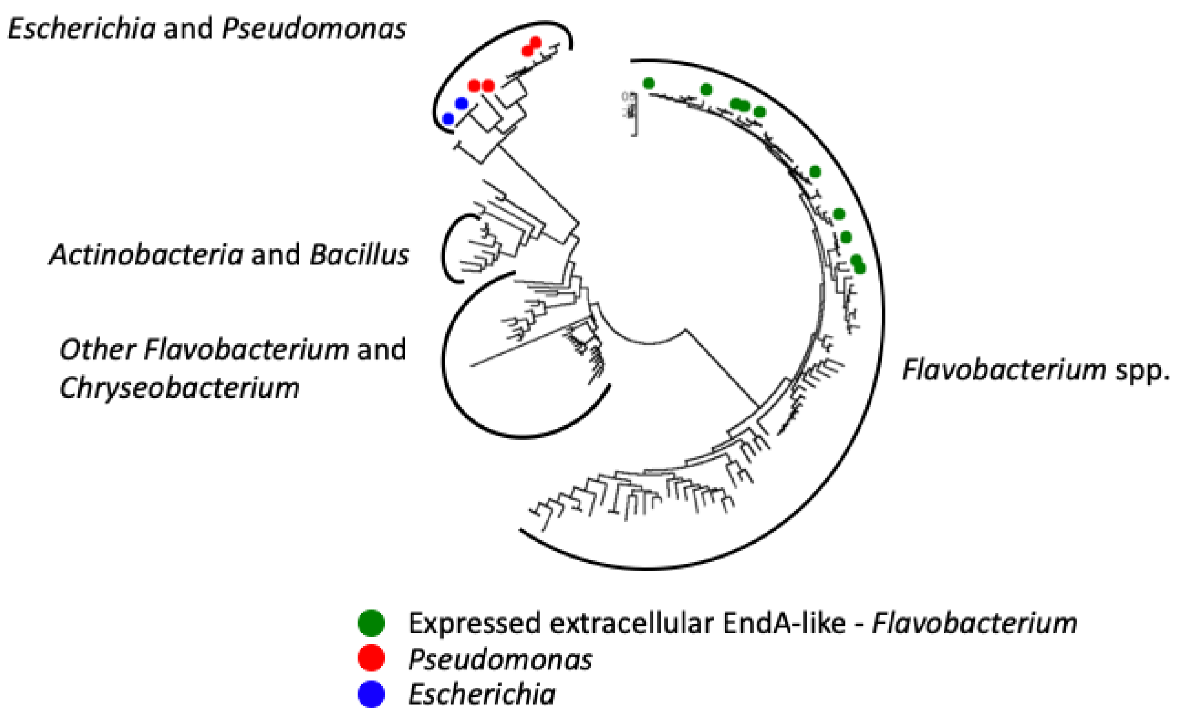


Figure. S4 Diversity of EndA-like sequences among soil/rhizophsere isolates including characterised proteins sequences form *Escherischia coli*. The evolutionary history was inferred by using the Maximum Likelihood method based on the JTT matrix-based model [1]. The tree with the highest log likelihood (-19815.95) is shown. Initial tree(s) for the heuristic search were obtained automatically by applying Neighbor-Join and BioNJ algorithms to a matrix of pairwise distances estimated using a JTT model, and then selecting the topology with superior log likelihood value. The tree is drawn to scale, with branch lengths measured in the number of substitutions per site. The analysis involved 151 amino acid sequences. All positions with less than 75% site coverage were eliminated. That is, fewer than 25% alignment gaps, missing data, and ambiguous bases were allowed at any position. There were a total of 194 positions in the final dataset. Evolutionary analyses were conducted in MEGA7 [2].

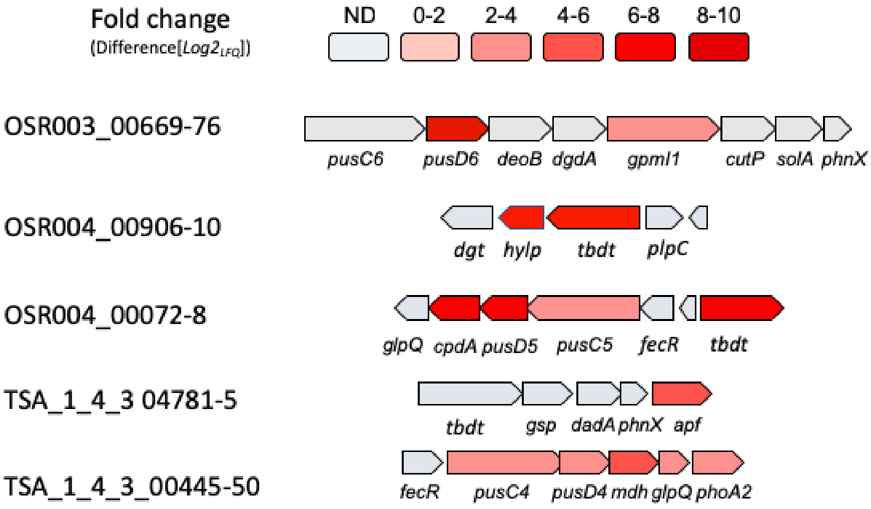


Figure S5. Genetic neighbourhoods of the proteins enriched in Pi-deplete exoproteomes of the various *Flavobacterium* isolates used in this study. Locus tags are given on the left-hand side. Results presented are the mean of triplicate cultures in each condition. The gene symbols are approximate annotations generated in PROKKA. In reality, the specific function of these proteins is unknown. Abbreviations: *GlpQ, glycerolphosphodiesterase; pusCD,* outermembrane transporter; *tbdt*, TonB-dependent outermembrane transporter; fecR, putative iron-responsive transcriptional regulator; *phoA*, alkaline phosphatase; *phnX*, putative phosphonatase; *dadA*, D-amino acid dehydrogenase; *cpdA*, 3',5'-cyclic adenosine monophosphate phosphodiesterase; *apf*, diadenosine tetraphosphate hydrolase; *mdh*, metal-dependent hydrolase; glucose permease; *deoB*, Phosphopentomutase; *dgdA*, 2,2-dialkylglycine decarboxylase; *gpmI1*, 2,3-bisphosphoglycerate-independent phosphoglycerate mutase; *solA*, N-methyl-L-tryptophan oxidase; *dgt*, deoxyguanosinetriphosphate triphosphohydrolase; *plpC*, S1/P1 nuclease/phospholipase C; *hylp*, hypothetical lipoprotein; *cutP*, carbohydrate/sugar phosphate permease.

High Pi Low Pi

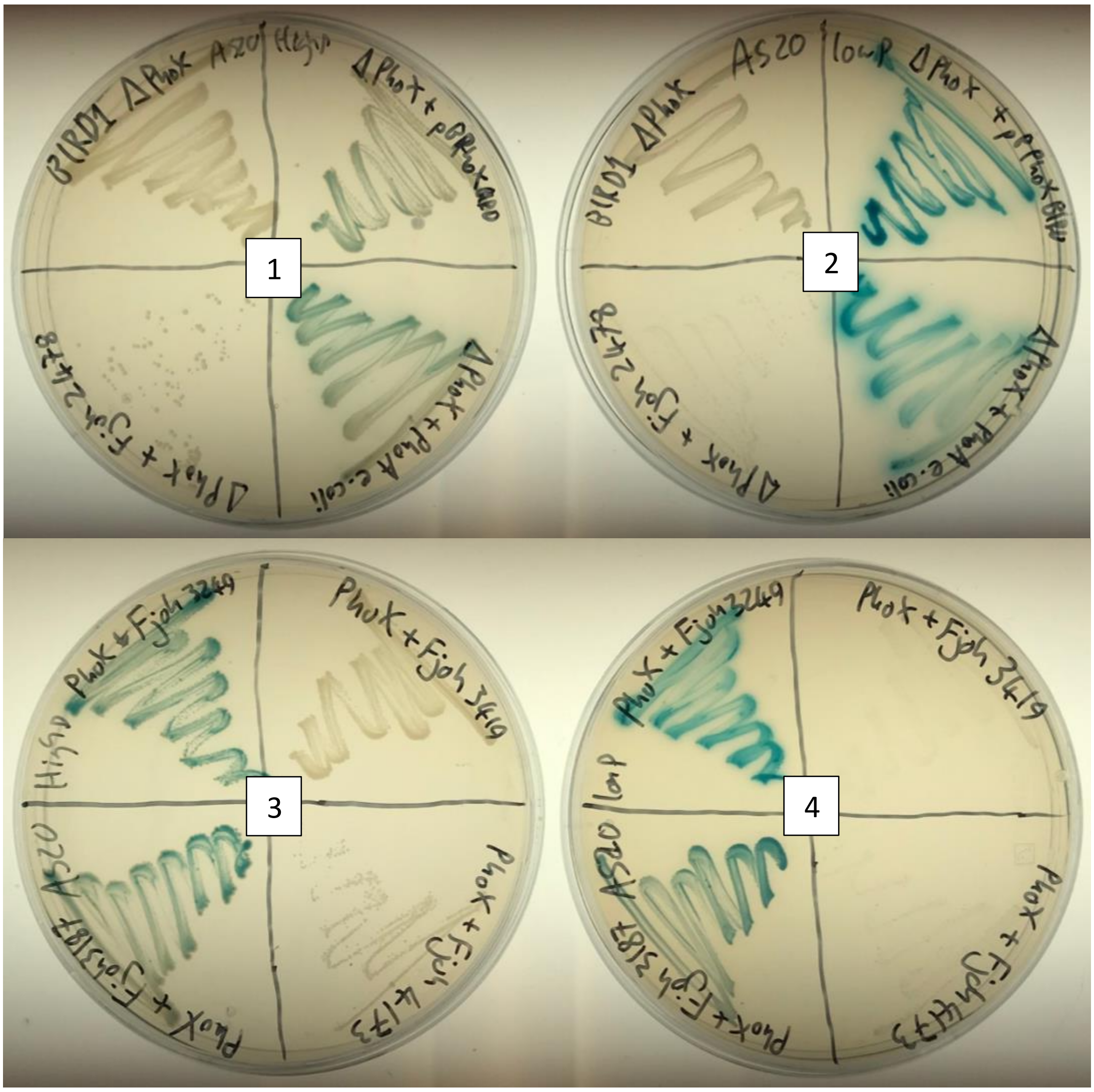


Figure S6. Alkaline phosphatase plate assay using the *Pseudomonas putida* BIRD-1 $\Delta$*phoX* mutant (top left, plates 1 & 2). 5-Bromo-4-chloro-3-indolyl phosphate (BCIP), more commonly known as XP, was used. The *phoX* mutant exhibited zero activity indicated by a lack of blue colouring which is created when XP is cleaved. Complementation with the native *P. putida* BIRD-1 restored the wild type phenotype. Heterologous expression of the two PhoA-like homologs (Fjoh_3187, bottom left P3 & 4 and Fjoh_3249, top left P3 & 4), also restored APase activity confirming their function. Neither Fjoh_3414 or Fjoh_4173 restored any phenotype. Interestingly, Fjoh_2478, the PhoX-like homolog, did not restore the phenotype. However, growth in this complemented mutant was inhibited which suggests that expression and subsequent export of the lipoprotein may be affected. Plates were left overnight at 30$^{o}$C.

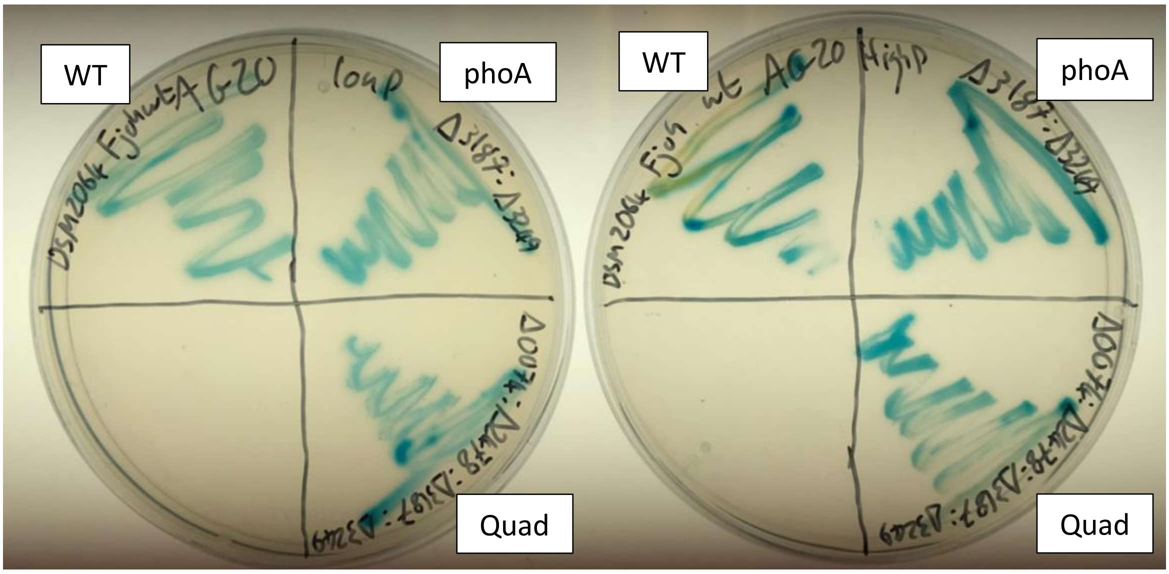


Figure S7. Alkaline phosphatase plate assay using the various *F. johnsoniae* APase mutants. Both the double *phoA* mutant and the quadruple mutant still displayed observable APase activity under Pi replete and Pi deplete growth conditions. (top left, plates 1 & 2). 5-Bromo-4-chloro-3-indolyl phosphate (BCIP), more commonly known as XP, was used. Abbreviations: WT, wild type; *phoA*, *ΔphoA1: ΔphoA2;* Quad, *ΔphoA1:ΔphoA2:ΔphoX:Δfjoh_0074*.

Table S14. List of primers used for mutagenesis in *Flavobacterium johnsoniae* DSM2064

| **Primer** | **Sequence 5'-3'** | **Function** |
|---|---|---|
| KO2064_3187_AF | GCAGCGGAAAAATTCGGGGGATCCTTGCATTCAGGTTTTCATTAGCG | For primer region A – PhoA2 |
| KO2064_3187_AR | TTCCGGGACCAAAGTTCCAGTGCTCATTCCGTC | Rev primer region A – PhoA2 |
| KO2064_3187_BF | TGAGCACTGGAACTTTGGTCCCGGAAGTGAAC | For primer region B – PhoA2 |
| KO2064_3187_BR | ATTACGCCAAGCTTGCATGCCTGCACGCAACAGATCAGGTTTG | Rev primer region B – PhoA2 |
| KO2064_3249_AF | AGCAGGGTTATGCAGCGGAAAAATTCGGGGACAATGCCAGCGATGACATC | For primer region A – PhoA1 |
| KO2064_3249_AR | CGGAACTGCTGTATGGATGTGTATTCCTGCGCC | Rev primer region A – PhoA1 |
| KO2064_3249_BF | GCAGGAATACACATCCATACAGCAGTTCCGGTTC | For primer region B – PhoA1 |
| KO2064_3249_BR | CTATGACCATGATTACGCCAAGCTTGCATGCTTTCCATCTGCCCAAAC | Rev primer region B – PhoA1 |
| KO2064_3250_AF | GCAGCGGAAAAATTCGGGGGATCCTGGATGCGGTTGACAAAGC | For primer region A – SusCD2 |
| KO2064_3250_AR | CAAGACCAGCACGTTTTGTCTAATGCCTGCTC | Rev primer region A – SusCD2 |
| KO2064_3250_BF | GCATTAGACAAAACGTGCTGGTCTTGGAGATC | For primer region B – SusCD2 |
| KO2064_3250_BR | ATTACGCCAAGCTTGCATGCCTGCAGGCCGGAGTAGCATCTGTAATATTTC | Rev primer region B – SusCD2 |
| KO2064_2478_AF | ATTACGCCAAGCTTGCATGCCTGCAGCCCTGCAGATATGATGG | For primer region A – PhoX |
| KO2064_2478_AR | GTATGAGGGTGAATGTGAAGTAAAATCGATGCTTG | Rev primer region A – PhoX |
| KO2064_2478_BF | ATTTTACTTCACATTCACCCTCATACCTGGC | For primer region B – PhoX |
| KO2064_2478_BR | GCAGCGGAAAAATTCGGGGGATCCTCGGTTTTGGAATAACTTGGC | Rev primer region B – PhoX |
| KO2064_4168_AF | GTTATGCAGCGGAAAAATTCGGGGGATCCTCAAAACTAAGCCAGCAGAC | For primer region A – SusCD1 |
| KO2064_4168_AR | CGTTAATGATGGTGTGAGAATCCGCCAAGAACTTC | Rev primer region A – SusCD1 |
| KO2064_4168_BF | TCTTGGCGGATTCTCACACCATCATTAACGTTCTC | For primer region B – SusCD1 |
| KO2064_4168_BR | CCATGATTACGCCAAGCTTGCATGCCTGCAACACTTCAACAGCTTCGTC | Rev primer region B – SusCD1 |
| KO2064_0074_AF | GTTATGCAGCGGAAAAATTCGGGGGATCCTAGAATACGAACCGGAAGC | For primer region A – 0074 |
| KO2064_0074_AR | TGAAGGTCTACAAATTCCATTTGAATCCTGACC | Rev primer region A – 0074 |
| KO2064_0074_BF | CAGGATTCAAATGGAATTTGTAGACCTTCACGCG | For primer region B – 0074 |
| KO2064_0074_BR | CCATGATTACGCCAAGCTTGCATGCCTGCATACCGGGGCTTTTGTTGG | Rev primer region B – 0074 |
